# Exploring the Conformational Landscape of Cryo-EM Using Energy-Aware Pathfinding Algorithm

**DOI:** 10.1101/2023.08.30.555633

**Authors:** Teng-Yu Lin, Szu-Chi Chung

## Abstract

Cryo-electron microscopy (cryo-EM) is a powerful technique for studying macromolecules and holds the potential for identifying kinetically preferred transition sequences between conformational states. Typically, these sequences are explored within two-dimensional energy landscapes. However, due to the complexity of biomolecules, representing conformational changes in two dimensions can be challenging. Recent advancements in reconstruction models have successfully extracted structural heterogeneity from cryo-EM images using higher-dimension latent space. Nonetheless, creating high-dimensional conformational landscapes in the latent space and then searching for preferred paths continues to be a formidable task. This study introduces an innovative framework for identifying preferred trajectories within high-dimensional conformational landscapes. Our method encompasses the search for the shortest path in the graph, where edge weights are determined based on the energy estimation at each node using local density. The effectiveness of this approach is demonstrated by identifying accurate transition states in both synthetic and real-world datasets featuring continuous conformational changes.

---

Cryo-electron microscopy (cryo-EM) entails imaging biological samples like viruses, proteins, and cells in a nearly-native frozen hydrated state at cryogenic temperatures using an electron microscope. Specifically, single-particle cryo-EM starts with images taken from each field of a frozen specimen containing target molecules in random orientations and positions. After computational analysis to estimate the particles’ orientations, the images can be classified and averaged, and a three-dimensional density map can be obtained via tomographic reconstruction [1].

This 3D reconstruction task is complex due to image distortion caused by the Contrast Transfer Function (CTF) and unknown projection angles with imperfect centering. To address these challenges, the homogeneous reconstruction scenario assumes that each recorded image *I*_1_, *I*_2_…, *I*_*n*_ captures a 2D projection of the same volume *V* at an unknown viewing angle, with an in-plane shift, as modeled by the following forward process:

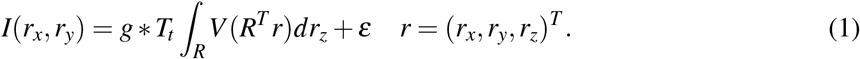

In this equation, *g* represents the point spread function (the inverse Fourier transform of the CTF), *R ∈ SO*(3) signifies the 3D rotation group representing the unknown orientation of the volume, *T*_*t*_ *∈* ℝ^2^ denotes the shift operator with an unknown in-plane shift 2D vector *t* and ε stands for noise. To recover the 3D density map, state-of-the-art algorithms use the Expectation-Maximization method to find the maximum a-posteriori probability estimate of the 3D volume *V*, while marginalizing over the posterior distribution of the unknown viewing angles ϕ = (*R,t*) and other latent variables, as expressed by:

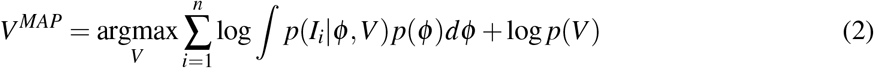

In practice, this method requires the generation of a low-resolution 3D initial model *V* ^0^ based on 2D class averages, common lines in Fourier space, or the stochastic gradient descent algorithm applied to Equation (2), and refining this model iteratively to achieve higher resolution.

Contrary to well-established methods like X-ray crystallography and Nuclear Magnetic Resonance (NMR), which predominantly determine single static macromolecular structures, single-particle cryo-EM has the capability to reconstruct 3D volumes for distinct groups of particles as classified by a 3D model. This methodology reveals diverse conformational states and elucidates the discrete structural heterogeneity inherent within a singular dataset. Such versatility positions single-particle cryo-EM as a potent tool to analyze heterogeneous datasets and determine the structures of macromolecular machines during dynamic processes.

Notably, recent advances in reconstruction models have effectively extracted structural heterogeneity on a continuous spectrum [3], thereby amplifying their capacity to manage intricate heterogeneous molecular structures. Among these techniques, neural network-based models [4, 5] have demonstrated proficiency in addressing the challenges of intricate heterogeneity reconstruction. In practical contexts, they produce high-resolution 3D volumes of primary states. Additionally, the latent spaces generated from these reconstruction models successfully capture the underlying conformational landscape, indicating their potential for subsequent dynamics analysis. The central premise is that the conformational movements of a protein can be represented as the paths of least energy on conformational landscapes. Given an initial and a final state, the trajectory bridging these states — which traverses the least cumulative energy — becomes pivotal in discerning the protein’s function. This can be achieved by employing pathfinding algorithms to explore the landscape.

Despite the promise of these advancements, the development and benchmarking of pathfinding algorithms in single-particle cryo-EM remains an emerging field. To bridge this gap, we first introduce a comprehensive synthetic generation workflow for continuous conformational changes, where dynamic transitions can be controlled using an occupancy map. Additionally, we propose innovative metrics for assessing the effectiveness of pathfinding algorithms. This benchmarking is vital not only for understanding the strengths and limitations of existing pathfinding methods but also for fostering the further development and validation of these techniques.

The other primary contribution of this work is that we propose an algorithm that navigates the typically high-dimensional latent space to find the most probable path. This diverges from traditional minimum energy path (MEP) algorithms, which typically search in a two-dimensional energy landscape ^1^. Instead, our approach directly constructs nearest neighbor graphs within the latent space — a method that is well-regarded for modeling relationships between data points in high-dimensional spaces. Additionally, we introduce a novel statistic that estimates local density at each node and assigns the graph edge weights as energy-aware values based on the Boltzmann factor. This approach streamlines the search for the minimum energy path in the graph using existing shortest path algorithms. Unlike traditional algorithms, our search process obviates the need to implement a 2D MEP algorithm by first embedding the latent space into a potentially limited-capacity, low-dimensional space. Finally, experimental results indicate that our proposed energy-aware weighting consistently exhibits a greater ability to identify transition sequences than conventional Euclidean distance weighting in both synthetic and real-world datasets.

## Results

### The proposed conformation analysis framework

Our proposed conformation analysis pipeline is depicted in Figure 1. To benchmark the methods and delve deeper into the impact of each component, we developed a simulation process in this study. This process can efficiently generate a conformational landscape in two or three dimensions using the atomic model and the designed occupancy map, as illustrated in Figure 1A. The particle stack is then generated based on the forward model (Equation (1)) and the occupancy map.

**Figure 1.**
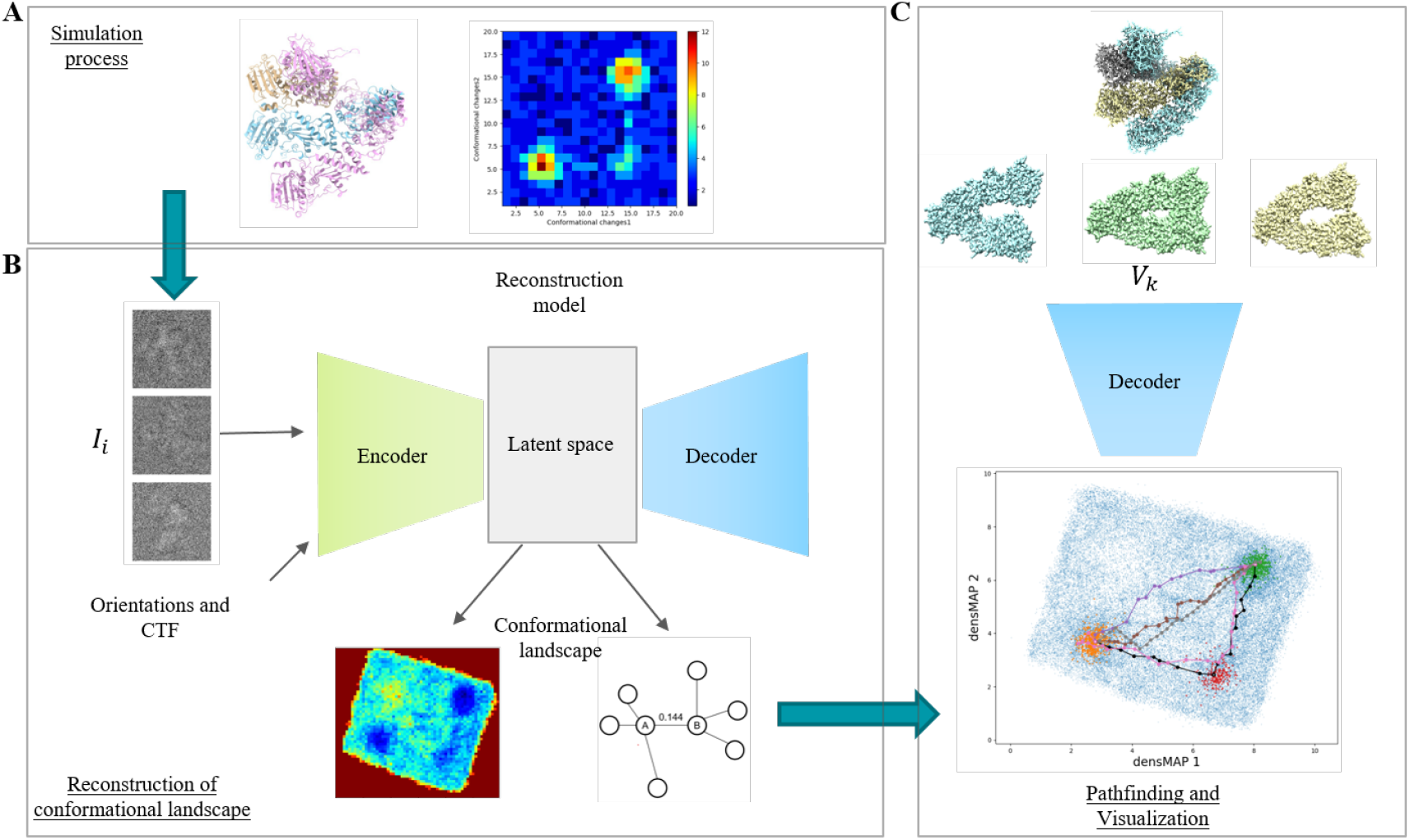
The proposed framework for conformation analysis in single-particle cryo-EM. **A**: To bench-mark the framework, a simulation process for continuous conformational changes is developed. This process generates a conformational landscape in two or three dimensions. Users have the flexibility to input different atomic models, design the corresponding occupancy map, and adjust the noise level to model different real-world scenarios. **B**: The reconstruction of the underlying conformational landscape is performed using a model, typically based on the auto-encoder framework. The particle stack, orientations, and CTF are input into the model for training. After the training, images are encoded into the low-dimensional latent space. The conformational landscape is derived using the two-dimensional energy landscape generated by the density-aware dimension reduction method or by constructing the nearest neighbor graph with energy-aware edges in the latent space. **C**: Once the conformational landscape has been obtained, several pathfinding algorithms are available to identify the most probable transition trajectory between different major states. Finally, the 3D density map representing different states can be generated by feeding the sampled points on the trajectories into the decoder.

To recover the underlying conformational landscape of the dataset, one can input either the synthetic dataset generated in Figure 1A or the dataset collected in real experiments into the reconstruction model. In the latter scenario, the orientations and CTF parameters will be determined by upstream software such as RELION [6] or cryoSPARC [7]. Typically, the reconstruction model is based on an auto-encoder framework [8–11] or eigen-analysis [12, 13], each identifying a low-dimensional (e.g., 8-10) latent representation of the underlying conformational landscape (Figure 1B, upper). In our framework, the conformational landscape is further transformed into different representations for downstream analysis. The first representation is a two-dimensional energy landscape created using the density-aware dimension reduction method and the 2D histogram (Figure 1B, lower-left). The second representation is the nearest neighbor graph with energy-aware edges, constructed in the latent space (Figure 1B, lower-right).

Once the conformational landscape has been recovered, the 2D MEP algorithm can be executed, given the starting and ending states. Alternatively, graph traversal algorithms can be performed on the graph to identify the preferred trajectories between the given states. The path found by these algorithms can be further visualized in two-dimensional space within our framework, again using the density-aware dimension reduction method (Figure 1C, bottom). Furthermore, the quality and faithfulness of the generated path can be examined using our proposed metrics. Finally, we can sample points along the trajectory or select points on the path with a higher local density, which may represent steady states. These sampled data points can then be fed into the trained decoder to generate 3D density maps representing different major states or a movie depicting the conformational changes in 3D space along the trajectories (Figure 1C, upper).

In this study, we choose the cryoDRGN [11] as our reconstruction model as it can recover the conformation landscape with enough precision (refer to Appendix B). In addition, we utilized the centers of the chosen major states in the latent space as inputs for the pathfinding algorithms. For the synthetic dataset, we treated the mean of data points, which belong to the same state in the latent space, as the centers. For the real dataset, we used the median of the data points to resist outliers. Furthermore, we obtained the ground truth trajectory by connecting the centers of the minor or transition states in the latent space. Our objective was to examine and compare three distinct pathfinding algorithms (refer to Methods) within our framework. These include the graph traversal algorithm from cryoDRGN, the search algorithm on a 2D energy landscape using POLARIS [14], and our novel energy-aware searching approach. We evaluated these algorithms on the two synthetic datasets and the two different latent spaces of the real-world EMPIAR-10076 [15] dataset. The aim was to find the intermediate transition states and generate trajectories that resemble the ground truth pathway. In the following experiments, we directly input the starting and ending centers for both the graph traversal algorithm and our energy-aware pathfinding algorithm. To search the 2D MEP using POLARIS, we constructed the energy landscape from the densMAP [16] 2D embedding space using a 2D histogram. Then, we used the corresponding starting and ending centers in the embedding space as inputs.

### Test on the Hsp90 dataset highlights the efficacy of the proposed framework in identifying transition states and the preferred trajectory

As a proof-of-concept, we apply our methodologies to the Hsp90 dataset, which exhibits two degrees of freedom in conformational changes (Supplementary Figure 4 and Supplementary Figure 5). The performance of the three pathfinding algorithms is shown in Figure 2. Notably, we plot the trajectories on the 2D embedding for visualization purposes, but the actual metrics computation occurs in the latent space, embodying the original conformational landscape recovered by cryoDRGN. We assessed the algorithms’ efficacy based on their precision in identifying the transition state and pinpointing the preferred trajectory.

**Figure 2.**
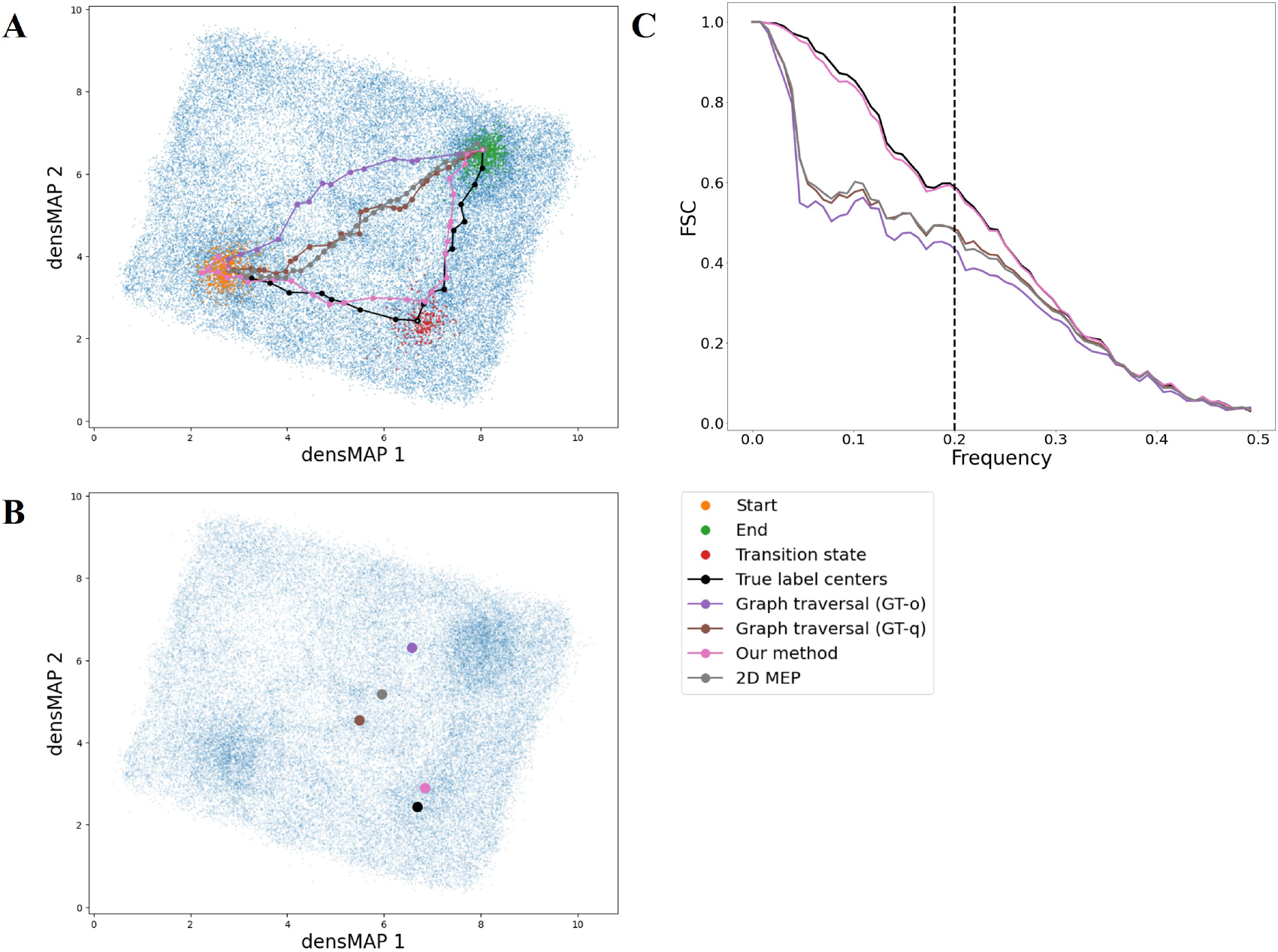
The results of the Hsp90 experiment. **A**: The paths identified by various algorithms are shown in the 2D visualization generated by the densMAP. Orange dots represent the starting state, while green dots denote the ending state. Red dots signify the transition state we aim to discover. The black line is the ground truth trajectory. Paths found by the different algorithms are color-coded: the default graph traversal (GT-o) path is purple, the path discovered through quantile searching threshold in graph traversal (GT-q) is brown, the path identified by searching on the 2D energy landscape using POLARIS (2D MEP) is gray, and the path found by our energy-aware pathfinding method is pink. **B**: This 2D visualization portrays the closest point for each path to the ground truth transition state center in latent space. The colors are consistent with those used in **A**. We utilize these nearest points to generate 3D volumes using the cryoDRGN decoder, which aids in calculating the Fourier Shell Correlation (FSC). **C**: The FSC curves between each algorithm’s volume and the ground truth transition state volume. The volume closest to the ground truth volume is used during the calculation for each path, with the color scheme corresponding to that in **A**.

As a first step, we computed the *FSC*_5Å_ (Equation (7)) between the volume generated by the nearest point on the predicted path and the actual 3D volume of the transition state used during the simulation. For reference, the volume formed by using the center of the ground truth label for the transition state in the latent space achieved *FSC*_5Å_ of 0.582 in the Fourier domain as shown in Table 1 and (Figure 2C). This reconfirms the accuracy of the recovered conformational landscape. Our energy-aware method closely followed this result, attaining a value of 0.578. On the other hand, the graph traversal method, which employs the quantile searching approach as a threshold (GT-q), gave a value of 0.480. When the POLARIS (2D MEP) method was applied to search on the 2D energy landscape, the value was 0.472. The default graph traversal (GT-o) method showed the worst performance, with the lowest value of 0.427. Furthermore, the *d*_Transition_(Equation (3)) metric shows that our predicted path was closest to the target transition state with a value of 0.897 in latent space (Figure 2B).

**Table 1:** Experimental results for metrics (Methods) across all datasets. The algorithms compared are Graph Traversal (GT-o), Graph Traversal with quantile threshold search (GT-q), 2D embedding with POLARIS (2D MEP), and our energy-aware pathfinding algorithm. GT-o was unable to identify a path in the fourth dataset. *g*_*i*_ signifies the state’s center *i* in the latent space, while *P* and *G* denote the sequence of points on the predicted path and the center points of the state on the ground truth path, respectively.

| **Hsp90** | **GT-o** | **GT-q** | **2D MEP** | **Our method** |
| --- | --- | --- | --- | --- |
| $FSC_{5\text{\AA}}(g_{Transition}, P)$ | 0.427 | 0.480 | 0.472 | 0.578 |
| $d_{Transition}(g_{Transition}, P)$ | 3.851 | 2.907 | 2.991 | 0.897 |
| $D_{ave}(G, P)$ | 2.478 | 1.789 | 1.957 | 0.628 |
| $D_{worst}(G, P)$ | 3.851 | 2.907 | 3.194 | 1.214 |
| **NLRP3** | **GT-o** | **GT-q** | **2D MEP** | **Our method** |
| $FSC_{5\text{\AA}}(g_{Transition}, P)$ | 0.254 | 0.283 | 0.403 | 0.485 |
| $d_{Transition}(g_{Transition}, P)$ | 3.689 | 2.915 | 3.265 | 0.602 |
| $D_{ave}(G, P)$ | 2.052 | 1.477 | 2.636 | 0.587 |
| $D_{worst}(G, P)$ | 3.689 | 2.915 | 6.696 | 1.154 |
| **EMPIAR-10076** | **GT-o** | **GT-q** | **2D MEP** | **Our method** |
| $FSC_{5\text{\AA}}(g_{C2}, P)$ | 0.887 | 0.969 | 0.919 | 0.942 |
| $FSC_{5\text{\AA}}(g_{E2}, P)$ | 0.905 | 0.907 | 0.925 | 0.993 |
| $FSC_{5\text{\AA}}(g_{E4}, P)$ | 0.923 | 0.913 | 0.807 | 0.989 |
| $d_{Transition}(g_{C2}, P)$ | 2.989 | 1.473 | 2.759 | 1.915 |
| $d_{Transition}(g_{E2}, P)$ | 2.835 | 2.967 | 2.177 | 0.769 |
| $d_{Transition}(g_{E4}, P)$ | 2.710 | 2.661 | 3.107 | 1.107 |
| $D_{ave}(G, P)$ | 2.936 | 2.502 | 3.922 | 1.162 |
| $D_{worst}(G, P)$ | 3.706 | 3.487 | 8.217 | 3.214 |
| **EMPIAR-10076 after analyze-landscape** | **GT-o** | **GT-q** | **MEP(2D)** | **Our method** |
| $FSC_{5\text{\AA}}(g_{C2}, P)$ | N/A | 0.852 | 0.933 | 0.894 |
| $FSC_{5\text{\AA}}(g_{E2}, P)$ | N/A | 0.904 | 0.938 | 0.962 |
| $FSC_{5\text{\AA}}(g_{E4}, P)$ | N/A | 0.817 | 0.917 | 0.932 |
| $d_{Transition}(g_{C2}, P)$ | N/A | 3.294 | 2.132 | 2.175 |
| $d_{Transition}(g_{E2}, P)$ | N/A | 3.388 | 2.505 | 1.787 |
| $d_{Transition}(g_{E4}, P)$ | N/A | 3.079 | 2.400 | 2.318 |
| $D_{ave}(G, P)$ | N/A | 2.972 | 3.884 | 2.427 |
| $D_{worst}(G, P)$ | N/A | 10.569 | 9.430 | 8.525 |

To quantify the overall similarity between the predicted path and the ground truth path, we calculate the average distances between the ground truth path and the predicted path using *D*_*ave*_ (Equation (4)). This yields a value of 0.628 for our energy-aware method. Furthermore, the largest discrepancy *D*_*worst*_ (Equation (5)) between the trajectories for our method and the ground truth is 1.214, considerably lower than the other methods.

In summary, these findings suggest that our proposed method outperforms the others in capturing the transition state and identifying the correct trajectory in the Hsp90 synthetic dataset. Finally, we can observe that the distance metrics are consistent with the densMAP visualization. The visualization presented in Figure 2A, as well as the metrics provided in Table 1, suggest that our path is the closest approximation to the ground truth, while the default graph traversal and 2D MEP perform the least effectively. Additionally, the graph traversal with quantile search demonstrates intermediate performance.

### Tests on NLPR3 datasets demonstrate the ability of our framework in identifying transition paths for molecules with more complex dynamics

We then examine the NLRP3 dataset in which the underlying molecule exhibits more complex dynamics (Supplementary Figure 6) and has a more intricate occupancy map than Hsp90 (Supplementary Figure 7). In this scenario, the three pathfinding algorithms were applied to the recovered conformational landscape with the aim of discovering the transition state and locating the preferred path, similar to the goal in the Hsp90 experiment.

In the NLRP3 experiment (Figure 3), the 3D volume from the center of the transition state achieves *FSC*_5Å_ of 0.497. Our energy-aware method attains a comparable correlation, with a value of 0.485, as shown in Table 1. The method of searching on the 2D energy landscape using POLARIS achieves a value of 0.403. The graph traversal with the quantile searching approach reaches a value of 0.283, while the default graph traversal method yields the lowest correlation of 0.254 (Figure 3C). Moreover, the *d*_Transition_ metric suggests that our predicted path was closest to the target ground truth transition state with a distance value of 0.602, making it the closest among all the algorithms.

**Figure 3.**
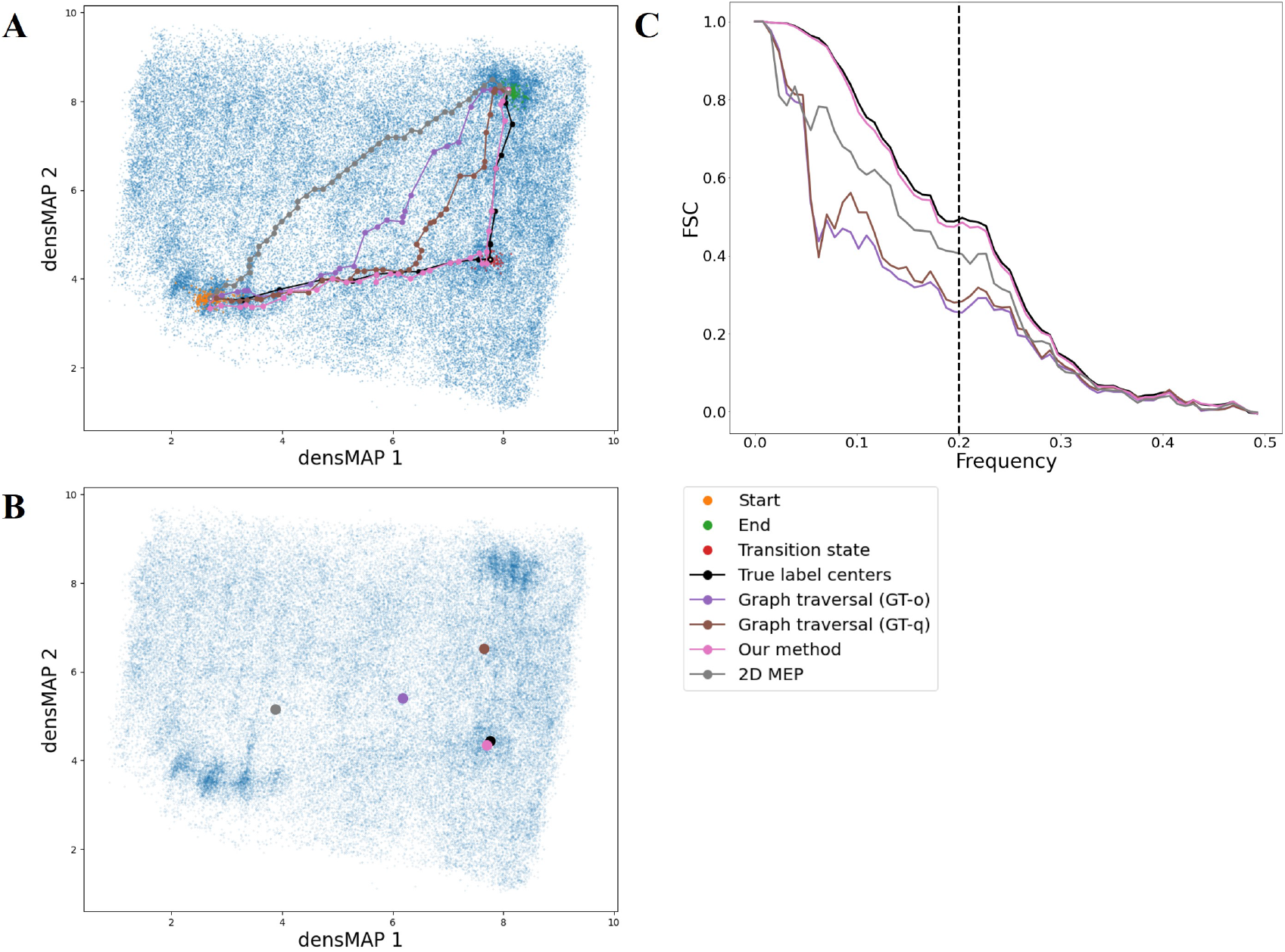
The results of the NLRP3 experiment. **A**: The 2D visualization, using the densMAP, depicts the paths identified by different algorithms. The orange dots signify the starting state, the green dots denote the ending state, and the red dots symbolize the transition state we aim to discover. The black line marks the ground truth pathway. The purple line corresponds to the path found by the default graph traversal (GT-o), the brown line is the path determined by the quantile searching threshold in graph traversal (GT-q), the gray line signifies the path searched on the 2D energy landscape using POLARIS (2D MEP), and the pink line represents the path found by our proposed energy-aware pathfinding method. **B**: The visualization shows the nearest point on each path to the ground truth transition state center in the latent space, with colors corresponding to those in **A**. We utilize these closest points to create 3D volumes using the cryoDRGN decoder, which assists us in calculating the FSC. **C**: The FSC curves between each algorithm’s volume and the ground truth transition state volume. The volume closest to the ground truth volume is used during the calculation for each path, with the color scheme corresponding to that in **A**.

On the other hand, the path located by our method is also closest to the ground truth path. The *D*_*ave*_ metric shows that the average distance is 0.587, and the largest discrepancy is 1.154 (Table 1) according to *D*_*worst*_. The graph traversal with quantile search ranks second, followed by the original graph traversal algorithm. In this case, the 2D MEP approach performs slightly worse with *D*_*ave*_ of 2.636 and *D*_*worst*_ of 6.696. Once again, the visualization (Figure 3A) and the distance metrics (Table 1) are in agreement. These results from a more complex setting align with those obtained from the Hsp90 experiment. Furthermore, they underscore the effectiveness of our framework in accurately capturing transition states and identifying the most probable path in biological molecules undergoing varying degrees of conformational changes.

### Our framework automatically reveals the intermediate state in the transition path of the bacterial ribosome dataset

Next, we evaluated the performance of the three pathfinding algorithms on the EMPIAR-10076 dataset, known for its significant compositional and conformational heterogeneity. In this study, our focus is on conformational heterogeneity. It’s important to note that the published labels suggest three potential conformational changes from the starting state B to the ending state E5 (Supplementary Figure 8). This complexity exceeds the straightforward pathway depicted in our synthetic dataset experiment. In the following experiment, we only provided the starting state B and ending state E5 as input to each algorithm. The remaining minor classes, which were previously discovered using several rounds of classifications, are then used as the target transition states that each algorithm seeks to discover. This approach allows us to evaluate the algorithm’s ability to identify potential transition states given the defined starting and ending points.

The search results for the path from the starting state B to the ending state E5 using the three different pathfinding algorithms are illustrated in Figure 4. Apart from our algorithm, the paths determined by the other algorithms appear to find a route that lies between two potential published paths. Consequently, we employed the metric *D*_ave_ to ascertain the closest published path and the path identified by each algorithm. This metric indicates that both the original graph traversal algorithm and the 2D MEP associate with the second path, denoted by the orange line in Supplementary Figure 8. In contrast, the other two algorithms align with the third path, depicted by the blue line. Notably, both published paths share transitions through states C2, E2, and E4. Such a similarity enables us to use these states as benchmarks to assess the effectiveness of the three pathfinding algorithms. Contrary to the scenario with the synthetic dataset, we don’t have a ground truth volume for comparison in this context. Nevertheless, when inspecting the volumes generated at the median point of these minor states, our method achieves the highest *FSC*_5Å_ values of 0.993 and 0.989 for the E2 and E4 states, respectively, as detailed in Figure 4C and Table 1. In contrast, the graph traversal algorithm employing the quantile selection strategy achieves the highest correlation for the C2 state at 0.969. Likewise, our methods record the smallest *d*_Transition_ values at E2 and E4, while graph traversal with quantile search has a lower *d*_Transition_ value at C2.

**Figure 4.**
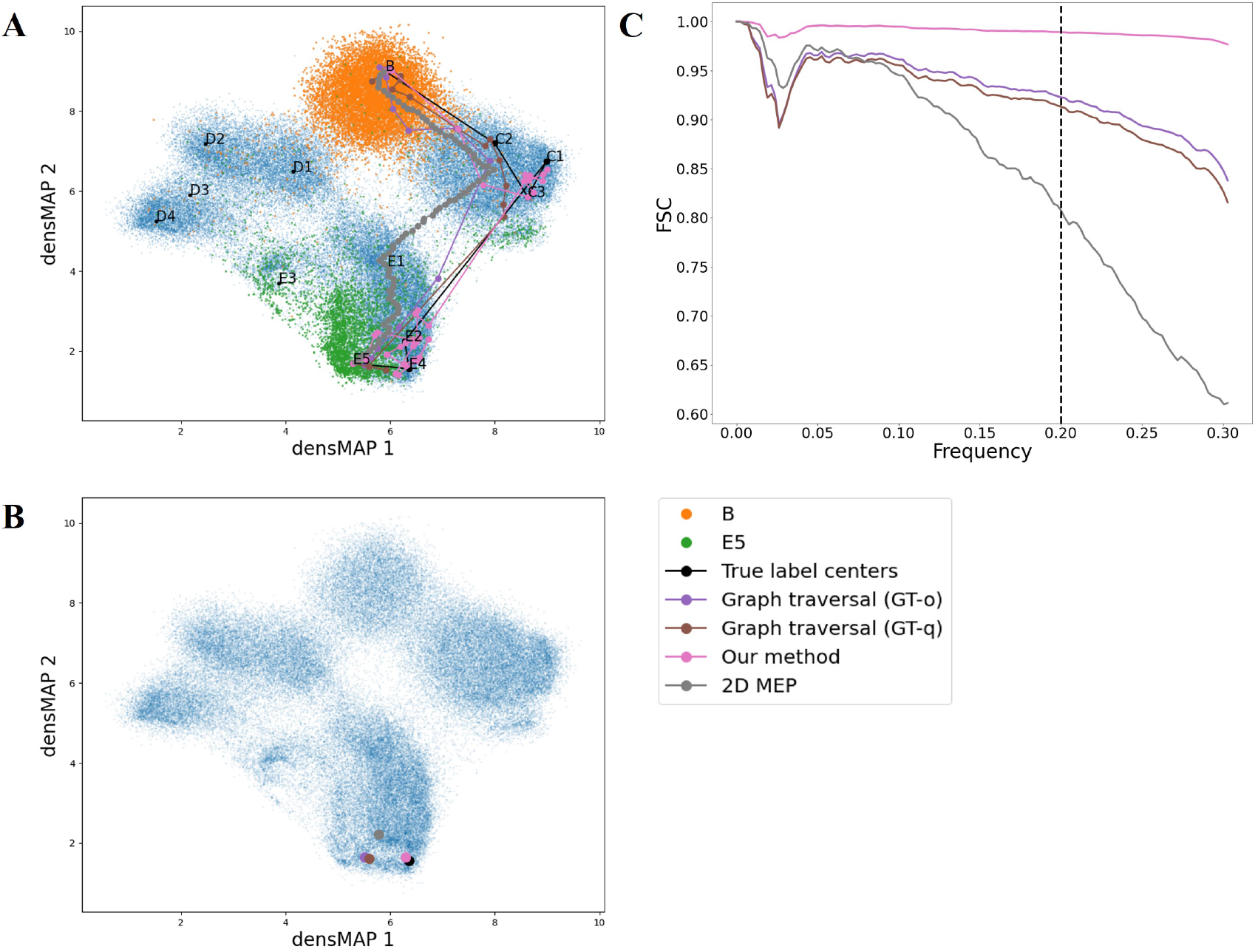
The results of the EMPIAR-10076 dataset. **A**: The 2D visualization, using the densMAP, illuminates the paths identified by the various algorithms. Orange dots represent the starting state, and green dots indicate the ending state. The black line marks the published path in the previous study. Colored lines represent paths identified by different algorithms: the purple line shows the path found by the default graph traversal method (GT-o); the brown line shows the path found by the graph traversal method using a quantile searching threshold (GT-q); the gray line shows the path found using POLARIS on the 2D energy landscape (2D MEP); and the pink line shows the path identified by our energy-aware pathfinding method. **B**: This visualization displays the closest point for each path to the ground truth label’s median point in state E4 in the latent space, with colors corresponding to those in **A**. We use these closest points to generate 3D volumes using the cryoDRGN decoder for FSC calculation. **C**: The FSC curves between each algorithm’s volume and the volume in state E4. The volume closest to the volume generated by the ground truth label’s center in the state E4 is used during the calculation for each path, with the color scheme corresponding to that in **A**.

To further examine the differences between each pathfinding algorithm, we compared their proximity to the ground truth trajectories. For a fair comparison, the path result derived from the original graph traversal algorithm and 2D MEP is compared to the second published path, the closest among the three published paths, and the other methods are compared to the third published path. Our method registers the lowest score of 1.162 in the average distance *D*_*ave*_ metric, indicating that our method generally finds the path closest to the ground truth center sequence. It also records the lowest score of 3.214 in the *D*_*worst*_ metric, signifying that it outperformed the other methods.

Finally, it is noted that our energy-aware pathfinding algorithm reveals the correct order of the sequence in the 2D embedding. The traversal order in the right-hand region is C2, C3, and C1, while the order in the bottom region is E2, E4, and E5. This suggests that our algorithm not only reveals the target intermediate states but also has the potential to determine the correct order of transition sequence while rejecting unfavorable transitions.

### Our framework is robust at searching the preferred trajectories after the analyze-landscape pipeline in the bacterial ribosome dataset

Given the inherent complexity of real-world datasets, the latent encodings that are learned might not always sufficiently capture structural heterogeneity. This shortfall can potentially result in suboptimal outcomes during the search phase of the pathfinding algorithm. To mitigate this limitation, we integrate a novel downstream analysis pipeline, termed analyze-landscape, into our framework as a post-processing step. This pipeline, as proposed by [17] (refer to Methods), primarily aims to map the latent space onto a more interpretable conformation space. It achieves this by capturing the heterogeneity in the output space formed by representative 3D volumes. We conducted experiments with this optimization to assess its impact on the pathfinding algorithm.

The resulting space is depicted in Figure 5. Notably, the minor states’ location on the 2D embedding aligns more coherently with the transition graph shown in Supplementary Figure 8. In particular, the ground truth trajectory no longer intersects itself in the C and E regions of the 2D space. Subsequently, we conducted a comparable experiment in this refined space, analogous to our approach with the original space, using the centers of states B and E5 as starting and ending points. The results are presented in Figure 5A. In this instance, the original graph traversal algorithm failed to find a path. Employing *D*_*ave*_ as the metric, the graph traversal with quantile search identified the second path. Conversely, the other methods highlighted the third path. This outcome justifies the necessity of performing a quantile hyperparameter search for the graph traversal algorithm, as recommended in our framework.

**Figure 5.**
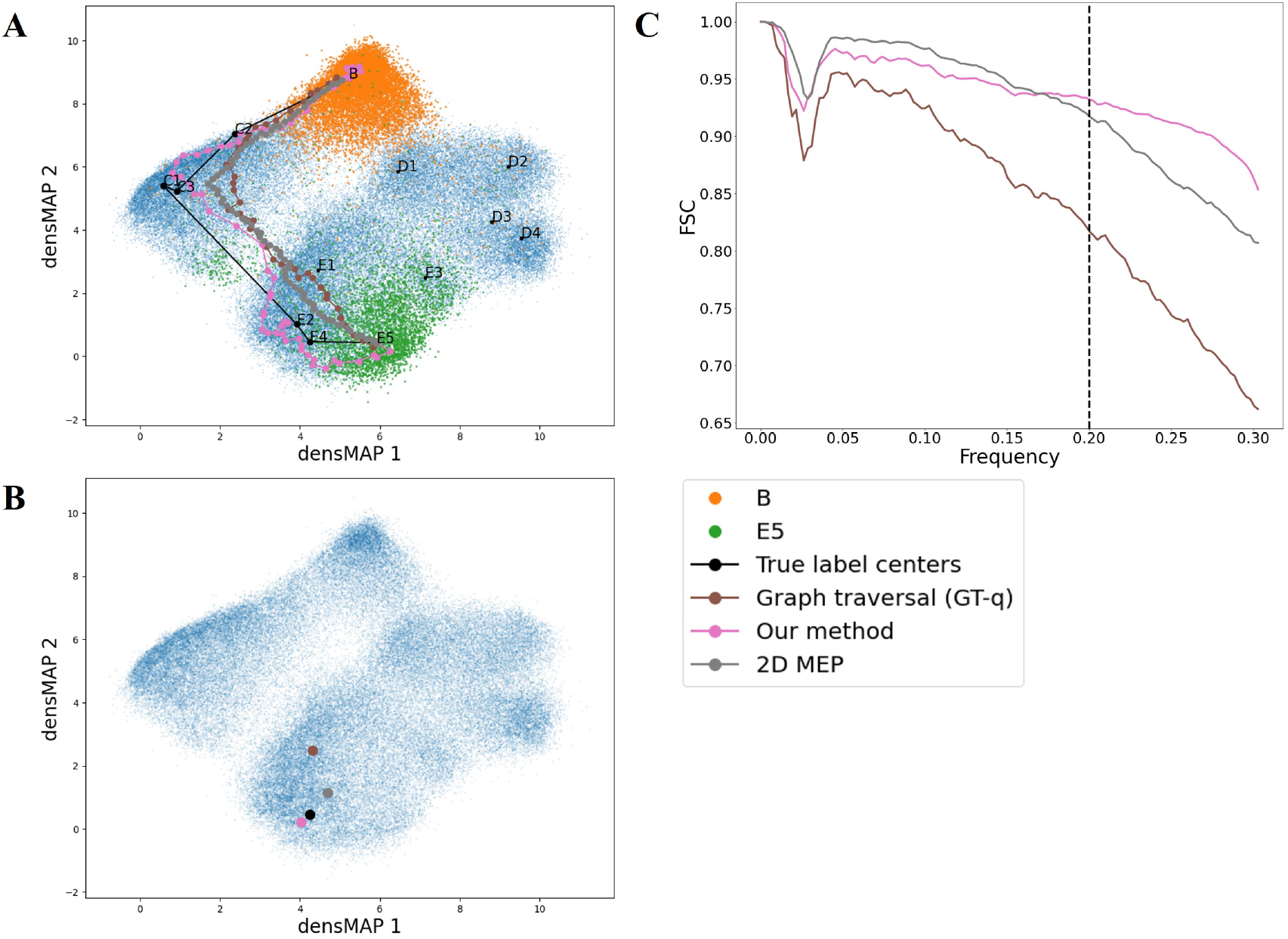
The results of the EMPIAR-10076 dataset after the analyze-landscape pipeline. **A**: The 2D visualization, created using the densMAP, emphasizes the paths identified by various algorithms. Orange dots signify the starting state, while green dots denote the ending state. The black line indicates the published trajectory in the prior research. Colored lines represent paths discovered by different algorithms: the brown line is the path found by the graph traversal method using a quantile searching threshold (GT-q); the gray line is the path determined using POLARIS on the 2D energy landscape (2D MEP); and the pink line is the path identified by our energy-aware pathfinding method. **B**: This 2D visualization illustrates the closest point for each path to the ground truth center of the state E4 in the latent space, with colors corresponding to those in **A**. We use these closest points to generate 3D volumes using the cryoDRGN decoder for FSC calculation. **C**: The FSC curves between each algorithm’s volume and the volume in state E4. The volume closest to the volume generated by the ground truth label’s center in the state E4 is used during the calculation for each path, with the color scheme corresponding to that in **A**.

For evaluative comparison, the shared states between the second and third paths were C2, E2, and E4. When calculating *FSC*_5Å_ relative to the volumes produced by the center of these minor states, our method yielded the top scores of 0.962 and 0.932 for the E2 and E4 states, respectively, as shown in Table 1. On the other hand, the 2D MEP presented the highest correlation for the C2 state, recording a value of 0.933. The *d*_Transition_ metric reflects a similar pattern, with the path identified by our methodology being the nearest to the E2 and E4 states. Furthermore, according to the overall distance metric, our method exhibited the closest proximity to the ground truth path, registering an average distance of 2.427 as represented by *D*_ave_. Additionally, it achieved a value of 8.525 in terms of *D*_worst_, illustrating that our energy-aware pathfinding method consistently outperforms the others in this new space. The results suggest that our proposed method is robust against changes in the underlying conformation space. Finally, it is noted that besides the original graph traversal algorithm, every other algorithm implemented in our framework identified a path proximate to the published route, as evident in Figure 5A. This emphasizes the strength of incorporating the analyze-landscape pipeline.

In conclusion, our framework pathfinding method has demonstrated the ability to find the transition and minor states we aim to recover according to the FSC and distance metric in the two synthetic datasets. This outcome gives us confidence that our method can accurately identify transition sequences. Moreover, our energy-aware pathfinding method successfully recognized the path closest to the ground truth trajectory, even while handling complex structural dynamics in real-world datasets.

## Discussion

This study primarily aimed to delve deeper into the downstream analysis of single-particle cryo-EM data, with a particular emphasis on identifying and validating the most probable transition paths. Our central focus was on developing an energy-aware pathfinding algorithm, which considers energy within the high-dimensional latent space to construct the most probable transition path among complex and heterogeneous biological macromolecules. To this end, we created synthetic datasets simulating continuous conformational changes. These datasets provided a controlled setting to validate and compare our proposed algorithm to others. We also designed metrics to benchmark the effectiveness of different pathfinding algorithms, offering a quantitative method to assess their ability to track conformational dynamics. This study also entailed designing a pipeline for implementing the POLARIS algorithm within a higher-dimensional space. This pipeline extended POLARIS’s ability to navigate 2D energy landscapes to higher-dimensional ones, using density-preserving manifold embedding techniques to maintain the integrity of the energy landscape. Finally, we also develop a quantile search procedure to enhance the performance of the graph traversal algorithm and show its advantage in the synthetic and real datasets.

Our method also demonstrated efficient execution times, completing search processes within a few minutes using default parameters for each experiment we discussed. Note that our algorithm’s efficiency surpasses that of the graph traversal with quantile threshold searching since the optimal threshold in our method is screened by a reasonable zero energy ratio controlling the space’s overall shape (see Methods). Consequently, we don’t need to perform actual searches on those suboptimal spaces. When compared to the strategy that searches on the 2D embedding using POLARIS, determining the optimal parameters for densMAP when building the 2D energy landscape can be time-consuming, thus prolonging the search process.

Overall, our energy-aware pathfinding algorithm demonstrated promising results, identifying the most probable transition paths in synthetic scenarios and showing potential for real-world dataset analysis. However, we acknowledge certain limitations associated with our approach. The first major limitation pertains to our assumption about the representation of structural heterogeneity within the latent encoding. Despite applying the cryoDRGN reconstruction model in our framework — a powerful tool in its own right — there exists the possibility that the latent space may not capture heterogeneity completely. As a result, using the latent space as a proxy may not always yield precise results. Specifically, unoccupied regions of the latent space may not necessarily correspond to highest-energy areas in the actual energy space. Yet, empirically, the latent space from cryoDRGN has demonstrated its effectiveness in analyzing structural heterogeneity in various real-world tasks. Therefore, using these representations as a foundation to approximate the true distribution of biomolecular structures is a sensible choice within our context. Furthermore, we tested a novel downstream analyze-landscape pipeline, which maps the latent space into a new space using 3D volume sketches. The performance of our method exhibited similarities across both original and new spaces, affirming the robustness of our framework.

The second limitation concerns the strategy of defining neighbors within a threshold, which might not provide an optimal estimation of occupancy and energy. This approach, similar to determining bins and counting points within each grid cell to construct a low-dimensional energy landscape for other minimum energy pathfinding algorithms, introduces ambiguity in delineating the precise shape of the energy landscape. This lack of clarity becomes especially pronounced without specific domain knowledge about the molecular structure under study, thereby affecting the selection of suitable bins for the low-dimensional energy landscape and the choice of the optimal threshold for our method.

Additional limitations stem from using Dijkstra’s Algorithm [18] to identify the minimum energy path in a graph. In situations where the graph contains a high number of zero-energy edge weights, our method may face difficulties. Consequently, the shortest path might inadvertently comprise many points from regions primarily characterized by zero edge weights, which is not always the desired outcome. This challenge led us to set the parameters for the ratio of zero energy to the total landscape at a low range by default, thereby limiting the prevalence of zero edge weights. However, this approach is tied to the shape of the landscape. We could artificially reduce the ratio further by opting for a smaller threshold, but our empirical findings indicate that such a strategy does not necessarily yield better performance.

To alleviate these limitations, future work lies in continuously evolving and enhancing our methods for constructing conformational landscapes from existing representation encodings. Enhancing the analyze-landscape pipeline [17] or extending the Manifold-EM [19] to a higher dimension are promising strategies. Such development could lead to advancements in our ultimate goal of identifying kinetically probable paths for biomolecules in single-particle cryo-EM. In conclusion, this study represents a significant step forward in the downstream analysis of single-particle cryo-EM data to understand protein functions, offering a framework for bench-marking pathfinding algorithms through a simulation process and innovative metrics. Moreover, this framework facilitates the identification of the most probable transition paths in complex biological macromolecules by employing different algorithms. The development and implementation of our energy-aware pathfinding algorithm demonstrate the feasibility of directly searching in high-dimensional latent spaces to pinpoint meaningful transition paths. While our method has its limitations, the positive results achieved using synthetic and real-world datasets provide a foundation for future research. Through continuous improvements and innovations in these areas, we can enhance our understanding of biomolecular dynamics, supporting breakthroughs in various fields of study, such as structural biology and bioinformatics.

## Methods

Benchmarking pathfinding algorithms in single-particle cryo-EM is a novel topic. To tackle this challenge, we first outline our synthetic process designed to simulate continuous conformational changes through occupancy maps. This workflow draws inspiration from recently published works on benchmarking various models [20, 21]. In addition, we propose novel metrics for comparing different pathfinding algorithms, with the aim of quantifying their ability to uncover crucial intermediate states and identify the correct traversing sequence. Secondly, we create a framework for implementing POLARIS in the latent space, utilizing a density-preserving manifold embedding technique to construct a 2D energy landscape. This density-preserving manifold embedding is also integrated into our analysis pipeline, proving to be an invaluable tool for visualizing various conformational states in a lower-dimensional space. Thirdly, we propose a quantile search procedure to improve the graph traversal algorithm. Finally, we introduce our fundamental concept of constructing a graph with energy-defined edge weights and present our energy-aware pathfinding algorithm.

### Simulation process for continuous conformational changes

There are five steps to synthesize datasets for our experiment (Supplementary Figure 9). Firstly, we generate a number of conformational states in atomic models, which exhibit two or three degrees of conformational changes in our experiment. We create these varying conformational states by applying rotations or movements to different chains of the atomic model, resulting in diverse structural variations that mimic the natural behavior of molecules. Secondly, we convert the atomic models into 3D density maps using the EMAN2 software [22], where we can control various parameters such as pixel size, resolution, and box size. This step ensures that the generated density maps accurately reflect the structural variations observed in the atomic models.

Thirdly, we apply an occupancy map to the 3D volumes generated in the second step. The occupancy map represents the number of clones in each conformational state, designed to emulate the single-particle cryo-EM image process where the microscope captures stable conformational states more frequently. To create the occupancy map (Supplementary Figure 10), we first generate data points in the same dimension as the degree of conformational changes we have chosen using a Gaussian mixture model. This approach allows the stable state to be represented as a group rather than a single state with numerous clones, resulting in a more continuous landscape that mimics the real energy landscape. We then divide the space into a number of segments corresponding to the conformational changes for each coordinate. Subsequently, we count the number of points inside each segment and rescale them into the desired occupancy levels to reduce computational costs.

In the fourth step, we apply random rotations and CTF to each clone volume, then project them into 2D images using the RELION software [6]. This step simulates the forward process (Equation (1)) of acquiring single-particle cryo-EM images with various orientations and imaging conditions. Lastly, we adjust the signal-to-noise ratio (SNR) to 0.1, approximating what is typically observed in realistic datasets. After stacking the 2D images, we use the image stack, ground truth viewing angles, and ground truth CTF as the input for training reconstruction models like cryoDRGN. This comprehensive synthetic data generation process enables us to create a robust and realistic dataset for evaluating the performance of our pathfinding algorithm.

### Proposed metrics for evaluating different pathfinding algorithms

Given that performance evaluation of pathfinding algorithms is a relatively new field, we propose two unique categories of metrics for assessment. The first category targets the alignment between the path generated by an algorithm and the ground truth in the latent space. The proximity can significantly influence its capacity to generate accurate volumes using the same decoder, thus providing a measure of the path’s precision and efficiency. We employ three distinct metrics to measure this closeness.

The primary objective of a pathfinding algorithm is to identify possible intermediate transition states given two major conformation states. Therefore, for each result produced by the algorithm, represented as *P* = *p*_1_, …*p*_*q*_, which is a sequence of data points in the latent space, we locate the point closest to the center point *g* of the transition state. This center point embodies the main transition state we aim to uncover, defined by the highest occupancy in the transition sequence for synthetic data or chosen in real-world datasets based on previous research. Furthermore, the center is calculated using the mean or median of all the data points that belong to the transition state. We then compute the Euclidean distance between this closest point and the transition center in the latent space. This distance serves as a measure of how accurately each algorithm can identify the transition states:

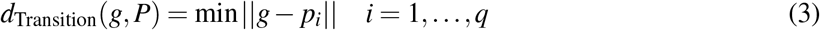

Here, ∥ *·* ∥ is the Euclidean norm. The second measure of closeness we utilize considers that the transition process typically consists of a sequence of intermediate states besides the important transition state and we denote the center of these states as a sequence *G* = *g*_1_, …, *g*_*k*_. For each state in this sequence, excluding the starting and ending states (which are given to the algorithm), we identify the nearest point to the state’s center on the algorithm-derived path. We then calculate the Euclidean distance between this point and the state in the latent space. Taking the average of these distances across all states in the sequence provides a summary measure of the average distance for the entire path:

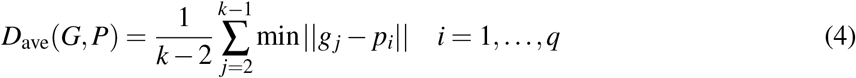

The third distance measure is the Hausdorff distance [23]. Given the ground truth center of state sequence *G* = *g*_1_, …, *g*_*k*_ and the algorithm’s prediction sequence *P* = *p*_1_, …, *p*_*q*_, the metric is defined as:

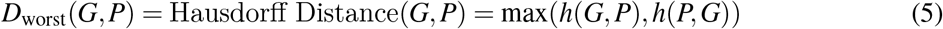

where

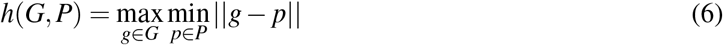

The Hausdorff distance between two sets of points is calculated by computing the Euclidean distance from a point in one set to the nearest point in the other and taking the maximum of these distances. This distance metric provides a worst-case scenario of the maximum discrepancy between two paths throughout the entire sequence. It reveals the maximum deviation between the sequence generated by a pathfinding algorithm and the ground truth path.

The second category of metrics relies on the volume generated using the decoder. We select the point closest to the center of the transition state in the latent space and employ the decoder to generate a 3D volume. We calculate the FSC for synthetic datasets with the ground truth volume used in the simulation process. For real-world data, where the ground truth volume is unavailable, the volume generated at the center of the transition state using the published label is employed. This measure assesses the correlation between volumes as a function of spherically-averaged radial shells in Fourier space:

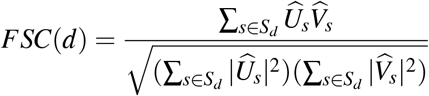

Where 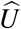 and 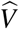 are the two volumes being compared, and *S*_*d*_ represents the set of voxels in a spherical shell at distance *d* from the origin. Although the original purpose of the FSC is to evaluate reconstruction performance through the consistency of two-fold cross-validation, we found that differences between results from various pathfinding algorithms can be captured at medium frequencies in the Fourier domain. Thus, we use a frequency of 0.2Å^*−*1^ (corresponding to a resolution of 5 Å) to compare the correlation with the ground truth volume. We define a metric that measures the FSC of volumes generated by the closest point on the predicted path *P* and the center of transition state *g* at 5 Å as:

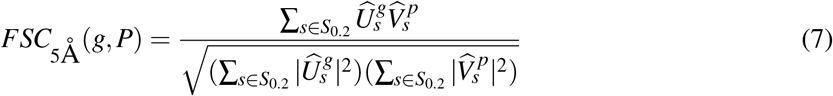

Here, 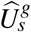 and 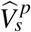 are the 3D volumes generated by the decoder at the closest point on the center *g* and path *P*, respectively. It is important to note that the resulting correlation may not provide robust evidence when the FSC is calculated without the ground truth volume. Therefore, we suggest interpreting the performance metrics in conjunction with the closeness metrics in the first category.

### 2D landscape visualization and POLARIS implementation using densMAP

To effectively visualize the high-dimensional structure heterogeneity representation and paths discovered by different pathfinding algorithms, we employ the density-preserving manifold embedding method, called densMAP [16], on the latent space. Although visualization tools like Uniform Manifold Approximation and Projection (UMAP) [24] and t-Distributed Stochastic Neighbor Embedding (t-SNE) [25] have been proven effective in both visualize heterogeneity [9, 11, 13] and building energy landscape [26] within single-particle cryo-EM data, they fail to account for the local density of data points in the original space. This makes them less effective for visualizing conformation mixtures, as data points that are closer together in the original space aren’t necessarily distinguished in the embedding space. Preserving the density structure is crucial when constructing an energy landscape using the histogram approach (Appendix A). Thus, we first discuss the densMAP visualization tool and then outline a pipeline for implementing POLARIS in the latent space using densMAP.

DensMAP generates embeddings that retain the density information of each point in the original space by utilizing a density measure called the local radius. This local radius is the average distance to the nearest neighbors of a point *x*_*i*_, denoted *R*_ρ_(*x*_*i*_). The local radius is defined as the expectation of the distance function over neighbors *x* _*j*_ with respect to a probability distribution ρ _*j*|*i*_:

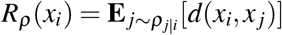

In line with the UMAP framework, the edge probabilities *P*_*i j*_ with points *x*_*i*_ in the original space are given as

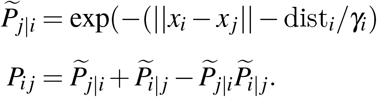

where γ_*i*_ is chosen adaptively and corresponds to the length-scale, and dist_*i*_ is the distance from *x*_*i*_ to its nearest neighbor. DensMAP renormalizes the edge probabilities *P*_*i j*_ to create a probability distribution for calculating the local radius in the original space, ρ _*j*|*i*_ = *P*_*i j*_*/* ∑ _*j*_ *P*_*i j*_:

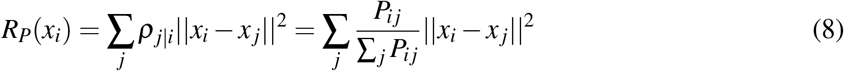

Following the same concept, the local radius for embeddings *R*_*Q*_ with points *y*_*i*_ in the embedding space is calculated based on the probability distribution ρ _*j*|*i*_ = *Q*_*i j*_*/* ∑ _*j*_ *Q*_*i j*_:

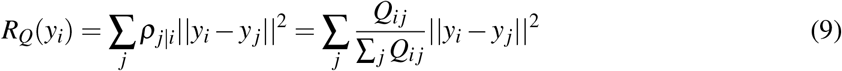

where *Q*_*i j*_ represents the edge probabilities computed in the embedding space:

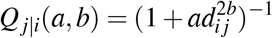

where *d*_*i j*_ represents the distance between *i* and *j* in the embedding, and *a* and *b* are additional shape parameters. Finally, densMAP modifies the loss function of UMAP by incorporating the correlation of the logarithms of the local radius in the original space and the embedding space. The densMAP loss function is optimized as follows:

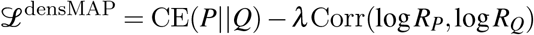

Here, CE represents cross-entropy, and λ is a user-chosen parameter that determines the importance of the density preservation term. Empirically, we found that using the densMAP visualization tool in single-particle cryo-EM provides a more accurate representation of the underlying data structure while maintaining the performance characteristics of UMAP (see Supplementary Figure 2). This finding underscores the potential of densMAP in visualizing the heterogeneity of single-particle cryo-EM data, mirroring its demonstrated capacity for visualizing scRNA-seq data in prior studies. In this study, the learning rate for densMAP is set to 0.001 for all the datasets, except that we use 0.005 for the dataset after landscape analysis. Additionally, `dens_frac` and `dens_lambda` are set to 0.7 and 3.0 in the two synthetic datasets, respectively, while all other parameters are left at their default values.

In principle, many conformational paths can connect a start conformation to an end conformation. However, most of these paths involve high-energy states, rarely populated under biologically relevant conditions. Due to the exponential nature of the Boltzmann inverse relationship between energy and occupation probability (Equation (11)), the lowest-energy conformational paths contribute maximally to function. Therefore, the search for the preferred path can be reformulated as a search for the MEP path. Considering the promising performance of POLARIS in identifying the most probable transitions in single-particle cryo-EM data within 2D energy landscapes (Appendix A), we aim to integrate POLARIS within the reconstruction model to assess its effectiveness in searching the 2D energy landscape. The integration process includes performing dimension reduction and building a 2D histogram on the embedding space as described below:

1. First, embed the latent space into a 2D embedding using densMAP.
2. Second, divide the 2D embedding space into a desired number of bins to create grids. Count the points within each grid cell to estimate their occupancies.
3. Third, construct a 2D energy landscape using the occupancies obtained in the previous step and the Boltzmann relation shown in Equation (11).
4. Finally, implement the POLARIS algorithm to search for the minimum energy path within the 2D energy landscape.

Different dimension reduction methods can alter the energy landscape upon which POLARIS performs its final search, making it essential to select an appropriate method for mapping the latent space into a 2D embedding space. To maintain consistency with the construction of the energy landscape, we choose to employ densMAP here. This strategy allows POLARIS to effectively identify the most probable transitions in the 2D embedding. While a density-preserving approach enhances the construction of the energy landscape, the performance of POLARIS remains sensitive to the quality of the embedding results (see Supplementary Figure 3). Moreover, determining the optimal parameters for densMAP introduces complexity to the implementation process. In this study, the energy landscape was constructed with 60 bins for each coordinate for the synthetic dataset and 100 bins for the EMPIAR-10076 dataset. In addition, the transition state weighting is turned on to avoid traversing over high-energy regions.

### Graph traversal and the quantile search algorithm

The first method involving direct traversal in the latent space is the graph traversal algorithm as described in [17]. This algorithm commences by creating a nearest neighbor graph from the latent encodings. A neighbor is defined if the Euclidean distance is below a threshold, computed from the statistics of all pairwise distances 2. The edges are subsequently pruned to ensure distant neighbors are disconnected from the given vertex. The Dijkstra algorithm is then deployed to find the shortest path connecting a series of points along the graph.

The method selects the median of all pairwise distances in the graph as the pruning point. Consequently, the resulting trajectory is less likely to pass through a low-occupancy region as distant neighbors are disconnected. However, it is evident that the threshold selection impacts the final generated trajectories. To address this, we implement a variant of the original method that performs a hyperparameter search on the threshold distance using quantiles. The best hyperparameter is chosen based on the statistic described in the subsequent section (Equation (10)). In the comparison of different pathfinding algorithms, we refer to the default version as GT-o and the variant as GT-q.

### Propose energy-aware pathfinding algorithm and search for the MEP in latent space

In this section, we present our innovative pathfinding algorithm, specifically designed to address the challenges associated with identifying the most energetically favorable transition pathways between molecular conformations directly from the latent space or other high-dimensional structural heterogeneity representations. Our approach consists of three main steps: constructing a graph, determining a threshold for occupancy estimation, and searching for the minimum energy path. Each step is explained in detail in the following paragraphs, highlighting our pathfinding algorithm’s unique features and advantages over existing approaches.

Firstly, we construct a nearest neighbor graph from the latent encodings by treating each data point in the latent space as a graph node. The neighbors are determined based on the Euclidean distance between the nodes. In our approach, the number of neighbors, *N*, is a parameter and influences the range of energy differences within the landscape, as the maximum occupancy is defined by the maximum number of neighbors for a node. For instance, if the graph is constructed with 50 neighbors, the maximum occupancy would be 50 and the minimum occupancy would be 0. This indicates that increasing the number of nearest neighbors generates more energy differences within the landscape. However, an increase in the number of nearest neighbors also escalates the computational cost and may not significantly alter the final results if the current differences already encompass the essential features of the landscape. Empirically, we found that *N* = 50 strikes a balance between computational cost and the adequacy of energy differences to identify an accurate path. Further increases in *N* do not affect the results much.

In the second step, we select a threshold to estimate the local density or the occupancies for each node in the constructed graph and define the edge weights between two nodes as an energy-like relationship (Supplementary Figure 11). Specifically, the threshold θ is a parameter that determines the size of the topological grids on the graph, similar to the grid used in the 2D histogram. For each node on the graph, we examine all its neighbors and compare their distances to the threshold. If the distance is greater than θ, it is considered a distant neighbor, potentially belonging to other states, and is therefore excluded from subsequent calculations. Following the pruning process based on the threshold, we count the number of remaining neighbors within the threshold to estimate the local density for each node. We found that using quantiles of the pairwise distances in the whole graph allows a systematic search for θ. Aligning with the concept of the energy landscape, we convert the occupancy estimations for each node into energy using Equation (11). The edge weights between two nodes on the graph are then defined as the average energy of these two nodes. This estimation can be seen as an efficient way to construct an *n*-dimensional histogram using the space defined by the nearest neighbor graph, offering several advantages. Firstly, this definition intuitively establishes a topological-like grid that performs a similar function to 2D grids. Secondly, it is more computationally efficient than other metrics like the local radius used in densMAP (Equation (8)) or the kernel density estimation. Lastly, it is robust against noisy points in the estimation since the threshold establishes a rejection bound to exclude outliers.

The threshold selection substantially impacts the pathfinding results, even more so than *N*. It modulates the overall shape and smoothness of the conformational landscape. For instance, selecting a smaller quantile as the threshold results in fewer neighbors remaining within the threshold for many nodes in the graph, engendering a landscape with many hills and fewer valleys. Conversely, opting for a larger quantile as the threshold increases the number of neighbors remaining within the threshold, yielding a more flattened landscape (Supplementary Figure 12). From a different perspective, this effect is analogous to constructing a 2D energy landscape with varying bins in each coordinate (Supplementary Figure 1). An increased number of grid cells in the 2D energy landscape is analogous to utilizing a smaller threshold to prune the graph, resulting in a higher likelihood of zero counts within the grids. This leads to a more pronounced separation of energy differences within the landscape, characterized by more distinct hills and fewer valleys. Following this concept, we can select an appropriate threshold by adjusting the ratio of zero energy in the entire landscape. To accurately depict the landscape, this ratio needs to be maintained within a specific range (*r*_*l*_, *r*_*u*_). This range serves as the final parameter for our algorithm and may require adjustment for different datasets. Empirically, we found a range between *r*_*l*_ = 0.01 and *r*_*u*_ = 0.1 to be effective for searching for the optimal quantile to use as the threshold. Finally, we utilize quantiles ranging from 0.1 to 0.9, with increments of 0.1 to search for the potential threshold.

In practice, the threshold that satisfies the desired proportion of zero energy in the entire landscape may not be unique. To address this, we calculate the average distance between each node and its neighbors along the path, then compute the mean of these average distances for each path that satisfies the constraint. For instance, if the nodes along the path are denoted *p*_1_, …, *p*_*q*_, and their neighbors *p* _*j*_, we calculate the mean of average distances for each node:

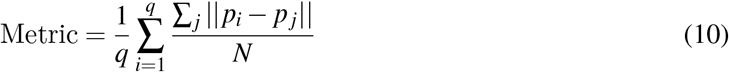

This metric assists in the final determination of the threshold, as nodes with lower energy should be closer to their neighbors.

Lastly, we use Dijkstra’s Algorithm to search for the minimum energy path. Given the non-negative edge weights defined based on average energy, Dijkstra’s Algorithm provides an efficient method for finding the shortest path within a specified graph. By estimating the local density and defining the edge weights as energy-like relationships, the shortest path corresponds to the minimum energy path. This path should represent the most energetically favorable transition pathway between molecular conformations. The summary of our algorithm is shown in Algorithm 1.

In summary, our pathfinding algorithm possesses several advantages. Firstly, it integrates energy information and searches directly within a high-dimensional latent space, thereby eliminating the need to select a dimension reduction method for constructing a low-dimensional energy landscape. Secondly, our algorithm’s parameters bear intuitive meanings. The parameter *N* determines the total number of energy levels in the landscape, while the parameter θ controls the overall shape of the landscape. Empirically, this intuitive understanding simplifies their determination and adjustment compared to the hyperparameter search of previous graph traversal algorithms, which depend on distance relations. In particular, performing the search with an appropriate zero energy ratio facilitates a more efficient implementation of our method, as opposed to selecting suitable densMAP and POLARIS parameters for searching in a 2D energy landscape or trying different parameter sets to boost the performance of graph traversal. Lastly, our algorithm’s application of energy-aware weights enhances the identification of a more probable trajectory compared to the graph traversal method in our experiment.

### Data preparation of the Hsp90 dataset

We generate a simulated dataset featuring two degrees of freedoms in conformational changes based on the Hsp90 molecule (PDB ID: 2CG9) [27]. The dual arm-like structures of this molecule allow us to conveniently depict two distinct directions of conformational changes. For each of the conformational changes, we create 20 states. The first direction pertains to the movement of one arm by 1 degrees for each state, and the second direction involves rotating the other arm by 2 degrees for each state (Supplementary Figure 4). These two directions correspond to the two coordinates in the occupancy maps. Using the synthetic process detailed in the previous subsection, we transform these conformational changes into 3D density maps. These maps, derived from atomic models, possess a resolution of 3 Å, a box size of 128, and a pixel size of 1 Å.

Subsequently, we devise an occupancy map with three groups of occupancies to simulate different conformational states of the Hsp90 molecule (Supplementary Figure 5). Two high-occupancy groups located in the lower-left and upper-right regions of the map, respectively, represent the initial and final states. Another group, with an occupancy that surpasses the background but is lower than the starting and ending states, is situated in the lower-right region of the map. This group signifies the main transition state that we aim to discover. To emulate the transition process and increase the chance of capturing more images during the conformational change, we also include a path connecting these three groups in the occupancy map. For a more realistic simulation, we introduce random noise into the background occupancy rather than using a constant 0 occupancy. This accounts for potential captures of conformations not part of the transition process in the dataset. The inclusion of random noise raises the complexity of the experiment, as pathfinding algorithms now have to differentiate between true transition states and random noise. After generating clones for each conformational state using the occupancy map, we construct 50 random viewing angles and project these clones into particle images. To achieve an SNR of 0.1, these images are convolved with a point spread function, and Gaussian noise is added. The resulting 56, 000 particle images constitute our dataset, which serves as the input for cryoDRGN.

### Data preparation of the NLRP3 dataset

For simulating more intricate conformational changes, we utilize the NLRP3 molecule (PDB ID: 6NPY) [28]. This molecule, which comprises a head, an arm, and a binding site ball adjacent to the arm, allows us to design three degrees of freedom in conformational change. For each of the conformational changes, we generate 20 states. The first change involves rotating the head by 2 degrees for each state, the second involves moving the binding ball by 2 degrees for each state, and the final one is represented by moving the arm by 2 degrees for each state, as shown in Supplementary Figure 6. Using the synthetic process detailed in the Methods section, we transform these conformational changes into 3D density maps. These maps, derived from atomic models, possess a resolution of 3 Å, a box size of 128, and a pixel size of 1 Å.

Following the methodology outlined in the simulation process, we create an occupancy map to simulate the transition process in the NLRP3 experiment. The occupancy map is designed similarly to the Hsp90 experiment, where the starting and ending groups have the highest occupancies, and a transition group has lower occupancy but is still higher than the background occupancy. We set 0 for the background occupancy and introduce 4, 000 random points as noise to mimic the real scenario. Additionally, to keep the computational requirements in check, we employ only 20 random viewing angles for generating the 2D images. Subsequently, these images are convolved with a point spread function, and Gaussian noise is added to reach a signal-to-noise ratio of 0.1. The final dataset, constituting 75,560 particle images, is then fed into cryoDRGN for further analysis.

### Data preparation of the EMPIAR-10076 dataset

To concentrate on the identification of potential transition states and simplify the training process, we adhere to the same workflow and use the intermediate results from the cryoDRGN team [29]. The box size is 256 pixel and the pixel size is 1.64 Å after downsampling. We use the published labels [15] as our ground truth to avoid reliance on clustering for analyzing structural heterogeneity. We remove states with labels A, F, and Discarded as they do not contribute to the conformational transitions from B to E5 (Supplementary Figure 13A) so that the remaining number of particles in the dataset is 87,461. Subsequently, we leverage the median points of B and E5 as the starting and ending points for each algorithm.

### Implementing the analyze-landscape pipeline for the EMPIAR-10076 dataset

We follow the cryoDRGN team’s three-step pipeline to map latent encodings into a more interpretable space represented by 3D volumes. The first step involves sketching the latent space. Given the high computational cost of generating a 3D volume for every point in the latent space, we simplify the latent space into a set of representative samples. This is achieved by using the centers of clusters identified by KMeans clustering, with the number of clusters set to 500. Subsequently, we generate 3D volumes with the cryoDRGN decoder using these selected points. The distribution of these representative 3D volumes represents the approximation of the population of the target protein.

The second step involves masking and dimension reduction. We apply masks to these volumes to filter out the background and focus on variations within the volumes. Subsequently, Principal Component Analysis (PCA) is applied directly to these masked volumes to reduce their dimensionality to a desirable level. The reduced dimensionality is set to 8, equivalent to the original latent space dimension.

The final step involves learning a mapping from the latent space to the newly constructed volume space. To accomplish this, we require more training data points. This is achieved by uniformly selecting additional 1000 points from the latent space. These points are used to generate additional volumes. We then apply the same mask created in the first step to these volumes and transform these 3D volumes into the same reduced dimensionality. A Multilayer Perceptron (MLP) network is subsequently trained to learn the mapping between the latent and volume spaces. Once the model is trained, it is used to predict the volume space of the entire dataset. The final volume space is used as the optimized latent space as shown in (Supplementary Figure 14).

### CryoDRGN training and generation of 3D volumes

For the training process on the two synthetic datasets, we follow the default architecture and training parameters as specified by cryoDRGN, albeit with a modification in the number of nodes in the hidden layers; this is adjusted to 256 to expedite the experimental process. Given that our objective is to focus on comparing the performance of different pathfinding algorithms, we feed the ground truth poses and CTF parameters into the reconstruction model. The generation of 3D volumes is conducted using the decoder of the trained cryoDRGN model. The Fourier Shell Correlation (FSC) is calculated using a custom script that can be accessed in the code repository.

## Data availability

The synthetic datasets were generated using custom code available at the following GitHub: https://github.com/tengyulin/synth_hsp90 and https://github.com/tengyulin/synth_nlrp3. The original EMPIAR-10076 dataset can be accessed at EMPIAR-10076. The trained cryoDRGN models, 2D embeddings, resulting paths and other experiment files have been deposited on Zenodo at https://zenodo.org/record/8229902.

## Code availability

The custom software framework and analysis scripts used in our study are available on the GitHub repository at https://github.com/tengyulin/energy_aware_pathfinding/tree/main and on Zenodo at https://zenodo.org/record/8229902.

## Acknowledgments

This work was supported by [MOST 110-2118-M-110-003-MY2] to S.-Z.C..

## Author contributions

S-C.C. conceived and designed the project. T.-Y.L. and S-C.C. developed algorithms. T.-Y.L. wrote the codes and conducted the analysis of the datasets. T.-Y.L. and S-C.C. analyzed the result and wrote the manuscript.

## A Background

### Neural network-based reconstruction model

The heterogeneity reconstruction problem is widely recognized as a primary challenge in single-particle cryo-EM. The standard approach to overcoming this challenge involves reconstructing *K* discrete volumes and generating *V*_1_, …,*V*_*K*_ with assignment probabilities π _*j*_, following a similar approach to the homogeneous problem (Equation (2)):

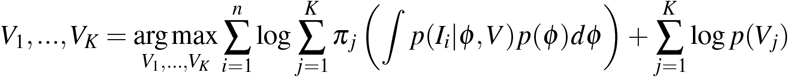

In practice, heterogeneity issues are often addressed through a hierarchical 3D classification process that involves refining subsets of the imaging dataset. The number of classes and initial models for refinement are manually selected. However, this method struggles when the structural heterogeneity comprises continuous conformational changes.

CryoDRGN (Deep Reconstructing Generative Networks) is one of the first neural network-based approaches for structural heterogeneity reconstruction in single-particle cryo-EM. It utilizes a variational autoencoder (VAE) framework to encode noisy 2D images with unknown viewing angles and imperfect centering into pose-invariant latent variables *z* that represent the structural heterogeneity. As per the VAE framework, the probabilistic encoder *q*_ξ_ (*z*|*I*) is a multilayer perceptron (MLP) with variational parameters ξ. When a 2D cryo-EM image *I* is inputted, the encoder outputs mean and variance, µ_*z*|*I*_ and Σ_*z*|*I*_ respectively, statistics that parameterize a normal distribution serving as the variational approximation to the true posterior *p*(*z*|*I*). The latent variable *z* is priorly set to a standard normal distribution *N*(0, I). Given latent variable *z* and Cartesian coordinates *k ∈* ℝ^3^ with positional coding, they are passed through the probabilistic decoder *p*_θ_ (*V* |*k, z*); the decoder outputs the slice at frequency *k* in Fourier space. Here, the coordinates are explicitly considered as the positions of each pixel in the 3D Fourier domain, thereby enforcing topological constraints between 2D projections in 3D through the Fourier slice theorem:

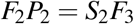

Here, *F*_*n*_ represents the Fourier transform in *n* dimensions, *P*_2_ is the projection operator from three to two dimensions, and *S*_2_ is the slice operator, extracting a 2D central slice perpendicular to the projection axis. In essence, each image captures an oriented central slice of the 3D volume in the Fourier domain.

To summarize, for a given input image *I*_*i*_, the encoder *q*_ξ_ generates two variational parameters, µ_*z*|*I*_*i* and Σ_*z*|*I*_*i*. Given the latent variables *z*_*i*_ *∼ N*(µ_*z*|*I*_*i*, Σ_*z*|*I*_*i*), and the rotation *R*_*i*_ for *I*_*i*_, the decoder *p*_θ_ then reconstructs the image pixel by pixel, taking the positional encoding into account. The microscope’s CTF and the phase shift *t*_*i*_ are then applied to the reconstructed pixel intensities. Following the standard VAE framework, the optimization objective function is the variational lower bound of the model evidence:

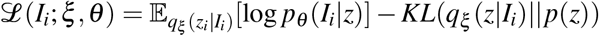

In this scenario, the expectation of the log-likelihood is estimated with a single Monte Carlo sample. This learning process functions as a denoising mechanism, preventing overfitting to noise from a single image, which could lead to increased reconstruction errors for other views. For further details on the rotation estimation, model architecture, and training process, the reader is referred to [11, 30]. In addition to producing high-resolution reconstruction results in real-world datasets, the latent variable effectively uncovers structural heterogeneity, paving the way for downstream analyses of structural dynamic processes based on the results from this powerful reconstruction model.

### Energy landscape

The energy landscape is a powerful tool for analyzing structural dynamics. This method is aligned with the fundamental postulate of statistical thermodynamics, which stipulates that in a thermodynamic system of fixed volume, composition, and temperature, all microstates with equivalent energy have equal probabilities [31]. Considering two microstates with corresponding energies *E*_1_ and *E*_2_. As the probabilities for these two microstates depend solely on their energies, we can denote *P*_1_ = *f* (*E*_1_) and *P*_2_ = *g*(*E*_2_), where *f* and *g* are functions of the energy whose forms are yet to be determined. Now, consider a third microstate which is a composite of microstates 1 and 2. The energy of this third microstate, denoted as *E*_3_, is the sum of *E*_1_ and *E*_2_. The probability of microstate 3 is yet another unknown function *h* of the energy: *P*_3_ = *h*(*E*_3_) = *h*(*E*_1_ + *E*_2_). Since the combined probability of simultaneous observation of microstates 1 and 2 equals the product of the probabilities of these two microstates, we can write:

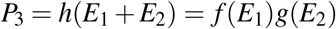

Taking the partial derivatives with respect to *E*_1_ and *E*_2_ on both sides, we obtain:

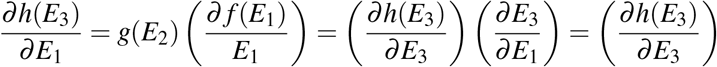

and

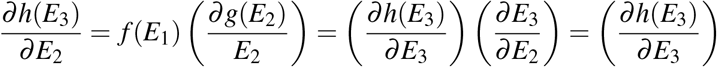

Since the right-hand side of both partial derivatives equals ∂ *h*(*E*_3_)*/*∂ *E*_3_, they must be identical. Furthermore, for the derivatives to be equal for distinct values *E*_1_ and *E*_2_, both sides must be constant. We denote this constant as *−*β:

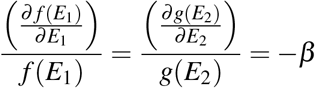

After solving these two differential equations, we find:

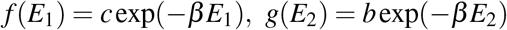

Here, *c* and *b* are constants. Generally, the probability of the *i*-th state in a system can be represented as:

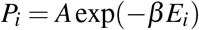

where *A* can be determined by ensuring the sum of probabilities of all possible states equals one:

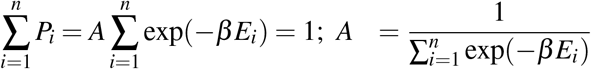

Finally, β can be deduced by studying a known system and generalizing the problem to β = 1*/K*_*B*_*T*, where *K*_*B*_ is the Boltzmann constant and *T* is the temperature. However, this aspect falls outside the scope of the current study.

In practice, we typically associate the major conformational state with the highest occupancy with the lowest energy, which is set to zero. Therefore, the ratio of the probability of any major conformational state to the lowest energy conformational state α is:

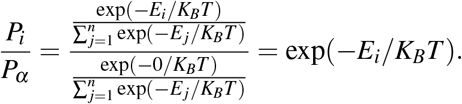

If the number of images in each conformational state is known, the probability ratio can be estimated by calculating the ratio of the number of images in each state. By taking the logarithm, we can derive the energy estimate [32]:

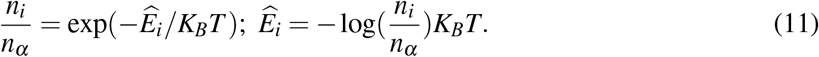

However, the number of conformational states in a given system is usually unknown for single-particle cryo-EM data. One potential approach is to devise an experiment that takes into account the known conformational change trajectory with timing information [32], effectively forcing the possible states to be observable through the microscope. Another common strategy involves selecting the reaction coordinates and dividing them into a set number of bins, each representing a conformation within that specific reaction coordinate. Usually, the number of reaction coordinates is chosen to be two or three, owing to the landscape’s complexity and the challenges of searching for the minimum energy path in a higher-dimensional space. The energy landscape, with its extensive hills and valleys, can then be navigated to identify a minimum energy path. This path represents the most probable sequence of conformational transitions.

**Supplementary Figure 1:**
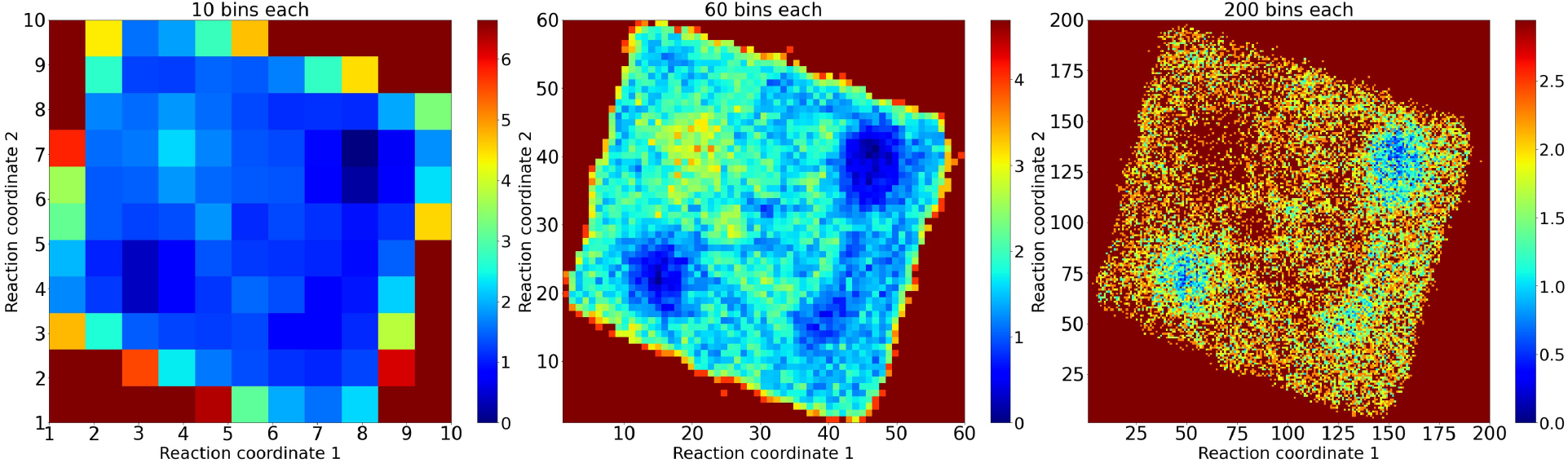
The impact of varying numbers of bins for constructing a 2D energy landscape, with the number of bins increasing from left to right. The color represents the energy of each grid cell. As the number of bins increases, many grid cells reach the maximum energy, resulting in a more distinct differentiation of energy levels within the landscape. Conversely, when a smaller number of bins is used, more grid cells exhibit zero energy, leading to a flatter landscape with fewer valleys.

### Pathfinding algorithms

In contrast to the reconstruction processes, the analysis of structural dynamics in single-particle cryo-EM is still evolving and remains relatively novel. Results from different models may necessitate entirely distinct analytical approaches. In this section, we outline various methods for identifying the most probable sequence of transition states occurring between any two conformational states.

As many reconstruction models depict structural heterogeneity as an *n*-dimensional manifold, the simplest and most intuitive approach involves sampling a pathway along the most variant coordinate. This can be accomplished by first using Principal Component Analysis (PCA) [33] or its variants [34, 35] to embed the heterogeneity representation coding, followed by sampling along the first or second component known as eigen-analysis. However, due to the complexity of biomolecular dynamics, this approach might not provide the most probable pathway. Moreover, sampling in only one or two directions may be insufficient to capture the true underlying conformational transitions.

The cryoDRGN team proposed an innovative method, known as graph traversal [11], for exploring trajectories within the latent space. This method can uncover more complex transition paths by searching in a higher-dimensional space. However, since the edge weights of the graph are defined as Euclidean distances, it may obtain suboptimal results when exploring conformational changes in biomolecules. Additionally, selecting the number of neighbors and determining the threshold can be challenging, as each choice can lead to vastly different paths. Another set of approaches involves searching for the minimum energy path within an energy landscape. The first approach is grounded in graph theory, where a graph is constructed and the most optimal path is identified based on a specified metric, such as the shortest path on the graph [36]. This method can be adapted to higher-dimensional energy landscapes [37]; however, constructing a high-dimensional energy landscape for cryo-EM datasets currently presents a challenge due to the lack of efficient methods, to the best of our knowledge. The second approach incorporates the string method [26, 38], which first connects two predetermined states with a straight line. This path is subsequently updated in a direction perpendicular to the gradient of the energy surface, continuing until the difference between successive updates falls beneath a user-determined tolerance. Nevertheless, this method may be affected by the selection of initial states and the determination of the tolerance value. Furthermore, its performance substantially degrades when searching within complex energy landscapes.

Lastly, the POLARIS (Path of Least Action Recursive Survey) algorithm seeks the minimum energy path in a 2D energy landscape using a recursive method [14]. It divides the landscape into equal 4^*n*^ blocks, determined by a user-defined parameter *n*, and pinpoints the minimum energy points in each block as candidates for the searching process. Another user-defined parameter, *r*, specifies the number of candidate transition points included in the search process. The search process recursively subdivides the potential path into pairs of points and searches based on these pairs with the same number of *r* until subdivision no longer reduces the energy. For instance, if the user-defined parameters are *n* = 1 and *r* = 1 with a starting point *S* and an ending point *E*, the possible path would involve searching from *S → C*_*i*_ *→ E* for four candidate transition points, where *i* = 1, …, 4. Subsequently, for the path *S → C*_1_, the algorithm would look for any other possible candidates to minimize the energy, thus refining the path further. This method allows a more thorough exploration of the energy landscape, as every point in the landscape can be a candidate when *n* is set to the maximum. Moreover, setting the parameter *r* = 1 can substantially reduce computation time, rendering the POLARIS method a promising option for searching minimum energy paths in 2D energy landscapes. However, computation time would increase dramatically when applying similar searching processes to higher-dimensional energy landscapes. Thus, this approach is extremely sensitive to the choice of a dimension reduction method capable of projecting higher-dimensional encodings to 2D and constructing an accurate 2D energy landscape. We will discuss this limitation further in the Appendix C.

## B The cryoDRGN can capture the conformation landscape with enough precision

In this section, we evaluate the capability of the current reconstruction model to extract conformational landscapes from cryo-EM images. According to a recent study [20], Manifold-EM [19] and cryoDRGN [11] are the two most promising methods for recovering the underlying conformational landscape. In this study, we chose cryoDRGN because it requires less hyperparameter tuning and can efficiently extract a conformational landscape with dimensionality larger than two 3. However, we considered a more realistic setting compared to [20]. Firstly, we assumed that the underlying degree of freedom of the conformational landscape is unknown, so we always learned an 8-dimensional latent space in all the experiments and used a density-preserving dimension reduction algorithm densMAP for visual assessment (see Methods). Additionally, we accounted for the CTF in the forward model (Equation (1)) and set the SNR to a lower value of 0.1 instead of 1 to make the synthetic dataset more closely resemble a realistic dataset.

We first examined the two synthetic datasets created for this study. The ground truth occupancy map of Hsp90 is designed to have two degrees of freedom, with two major states located in the lower-left and upper-right corners, respectively. These states show a high occupancy level, as demonstrated in Supplementary Figure 5. A transition state is added in the lower-right corner with higher occupancy than the background but less occupied than the major states. For the NLPR3 dataset, we considered a more complex conformational dynamic than in the Hsp90 experiment. The occupancy map was similarly designed but had three degrees of freedom as illustrated in Supplementary Figure 7. The starting and ending states, located in the corner of the 3D box, have the highest occupancies. A transition group in another corner exhibits slightly lower occupancy, though it is still higher than the background occupancy. The noise was added to the background to mimic the real-world scenario for both datasets. For the EMPIAR-10076 dataset, we do not have the ground truth occupancy map, but we do have the published labels for each major and minor state as well as the transition path (Supplementary Figure 8) released by a previous study [15] (see Methods for more about the data preparation).

The conformational landscapes of the two synthetic datasets, as recovered by cryoDRGN, can be visualized in Figure 2 and Figure 3 using the densMAP. It’s clear that cryoDRGN successfully captures the two major states and transition states at the corner, all of which have a higher occupancy than the background. In addition, the major states enjoy a higher occupancy level than the transition state. For the EMPIAR-10076 dataset, the recovered landscape is illustrated in Supplementary Figure 13, and the optimized conformational landscape by the analyze-landscape pipeline [17] is depicted in Supplementary Figure 14. We assigned different colors to data points that belong to different classes according to the label. Clearly, the conformational landscape successfully separates each major and minor state. In addition, states that are closer in the plot of the transitional path (Supplementary Figure 8) also appear closer on the scatter plot. From these observations, we infer that cryoDRGN can learn a reasonably accurate conformational landscape from both synthetic and real datasets.

## C Examining the impact of 2D embedding on POLARIS

In this section, we discuss the details and constraints of searching on a 2D energy landscape using POLARIS. As noted in the Methods, we opted for a density-preserving manifold embedding technique over commonly used approaches such as UMAP or t-SNE. This choice is critical when using the bin-counts approach to construct an energy landscape (Appendix A). To demonstrate its significance, we compared embeddings using UMAP and densMAP in the Hsp90 experiment (Supplementary Figure 2). After defining the starting, ending, and transition state centers as inputs, our objective was to identify a path akin to the black line, symbolizing the actual center of the transition sequence. Empirically, we found that the low-energy (high-density) region using UMAP differs from the one using densMAP and may distort the energy landscape. This difference influences the result of minimum energy pathfinding by POLARIS, leading to the predicted pathway deviating from the true sequence.

The other factor that may impact the final selected path is the relative position of the start, end, and transition center on the 2D embedding. To illustrate this impact, we used the same occupancy map to create a new dataset in which the first conformational change is reduced to 0.5 degrees for each state, and the second conformational change is reduced to 1.44 degrees for each state. This new dataset was created following the same workflow as in the Hsp90 experiment, and we used the same training processes to obtain the latent space. Subsequently, we constructed a 2D energy landscape using the same bin size and applied the same parameters in POLARIS to find the MEP. The results are presented in Supplementary Figure 3.

When comparing with the true sequence, we observed that even though the energy landscape looks similar (Supplementary Figure 3A and Supplementary Figure 3C), with three regions corresponding to the starting, ending, and transition states, the outcomes are completely different. The new dataset accurately reveals the path. This difference can be attributed to the fact that, in these cases, the regions within the start, end, and transition areas have roughly the same energy. Consequently, if no significant low-energy regions are present to explore, the MEP would depend more on the length of the distance between these states. In the dataset with lesser rotation and movement, the distance between the starting and ending points is 25.239, which is larger than the original dataset’s 24.352, and the distance passing through the transition state is 61.497 smaller than the original dataset’s 65.676. Therefore, POLARIS explores the transition state region instead of traversing over the middle region for the dataset with lesser rotation and movement. These effects substantially impact performance and introduce uncertainty when using POLARIS to search on a 2D energy landscape.

**Supplementary Figure 2:**
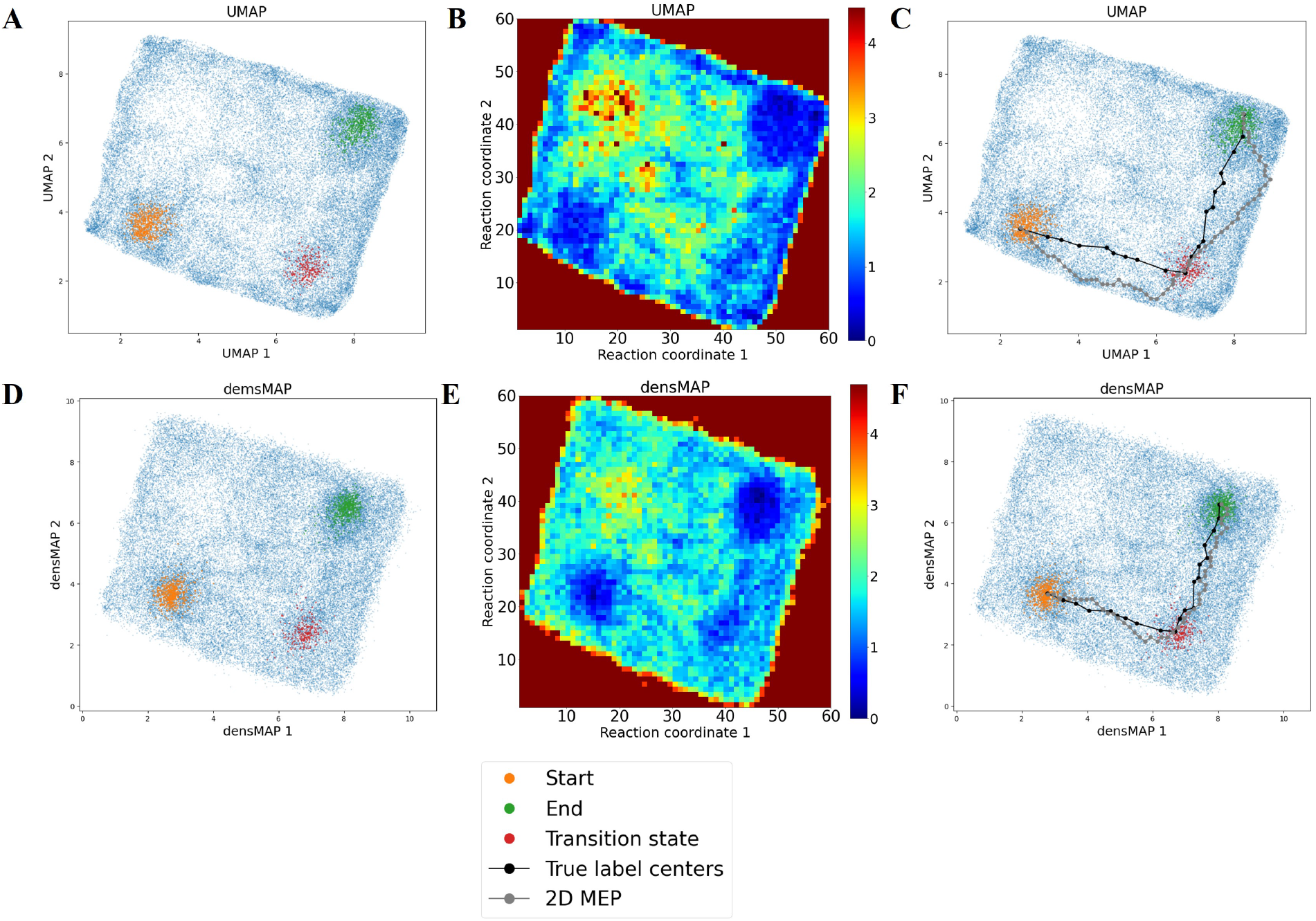
Comparison of the impact of embedding selection when using POLARIS to search for the MEP on a 2D energy landscape. **A** displays a 2D visualization after performing UMAP on the Hsp90 experiment latent space. Orange dots represent the starting state, while green dots denote the ending state. Red dots signify the transition state. **B** shows the energy landscape constructed by a 2D histogram after selecting 60 bins for each coordinate, and **C** contrasts the path identified by POLARIS (depicted by the gray line) with the ground truth transition sequence (represented by the black line). Figures **D, E**, and **F** represent results obtained using densMAP, corresponding to the same types of visualizations as in **A, B**, and **C**. In the UMAP scenario, the low-energy region is shifted towards the landscape’s edge, which inaccurately represents the transition sequence’s center. Conversely, when employing densMAP, the low-energy regions accurately align with the ground truth center sequence. Note that in this experiment, the transition state center is also provided to the algorithm in addition to the start and end state.

**Supplementary Figure 3:**
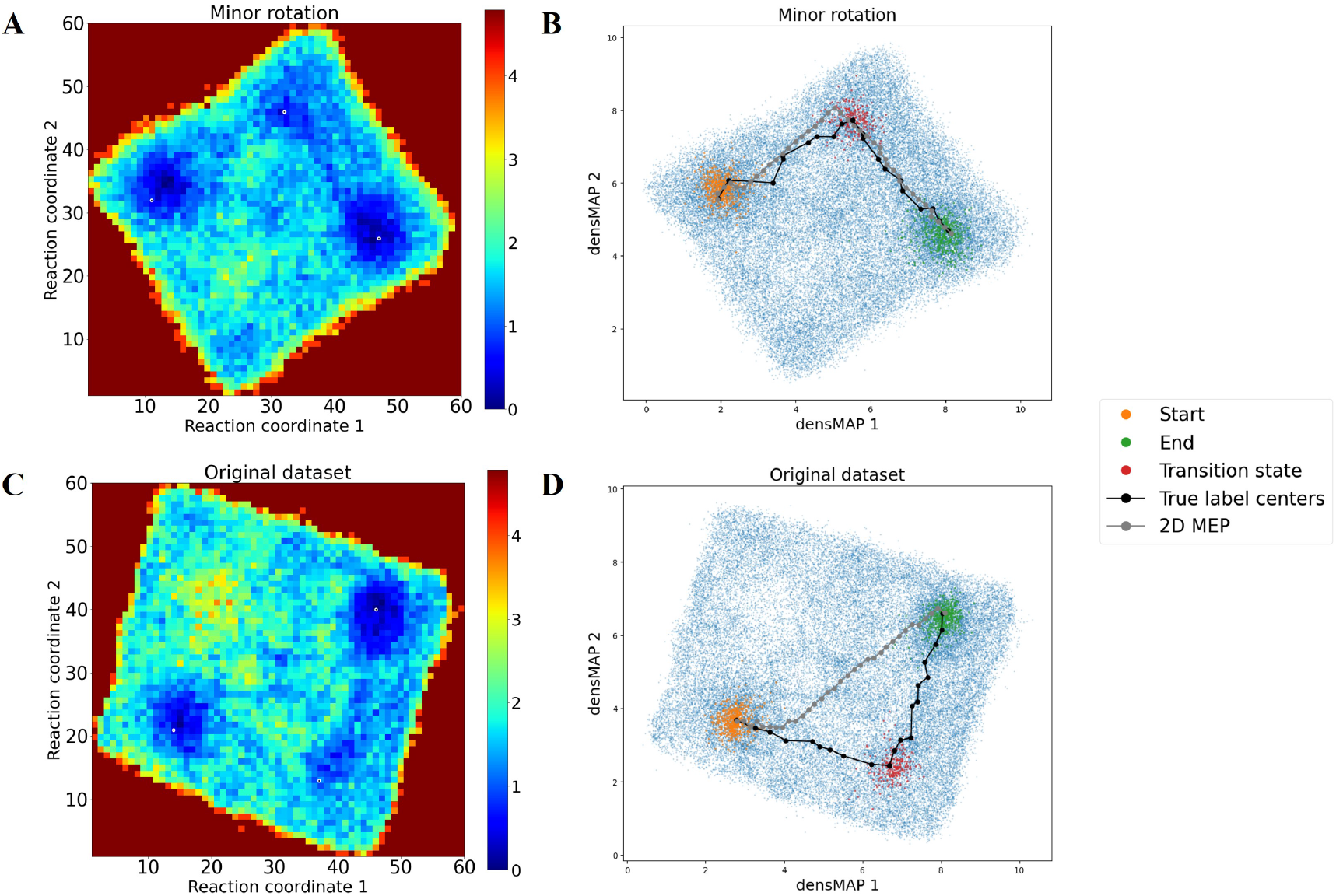
Comparison of the new dataset, with less rotation and movement, when searching for the MEP using POLARIS. **A** and **B** represent results from the new dataset, in which the first conformational changes are reduced to 0.5 degrees each, and the second conformational changes are reduced to 1.44 degrees each for the Hsp90 molecule. **A** shows the 2D energy landscape constructed after embedding using densMAP, with 60 bins selected for each coordinate. **B** contrasts the predicted path (depicted by the gray line) with the ground truth center sequence (represented by the black line). **C** and **D** depict the results from the original Hsp90 experiment, using the same representations as in **A** and **B**. Although the two energy landscapes appear highly similar, the paths found by POLARIS are entirely different. The predictions on the dataset with less rotation and movement accurately capture the transition state region.

## D Supplementary Figures

**Supplementary Figure 4:**
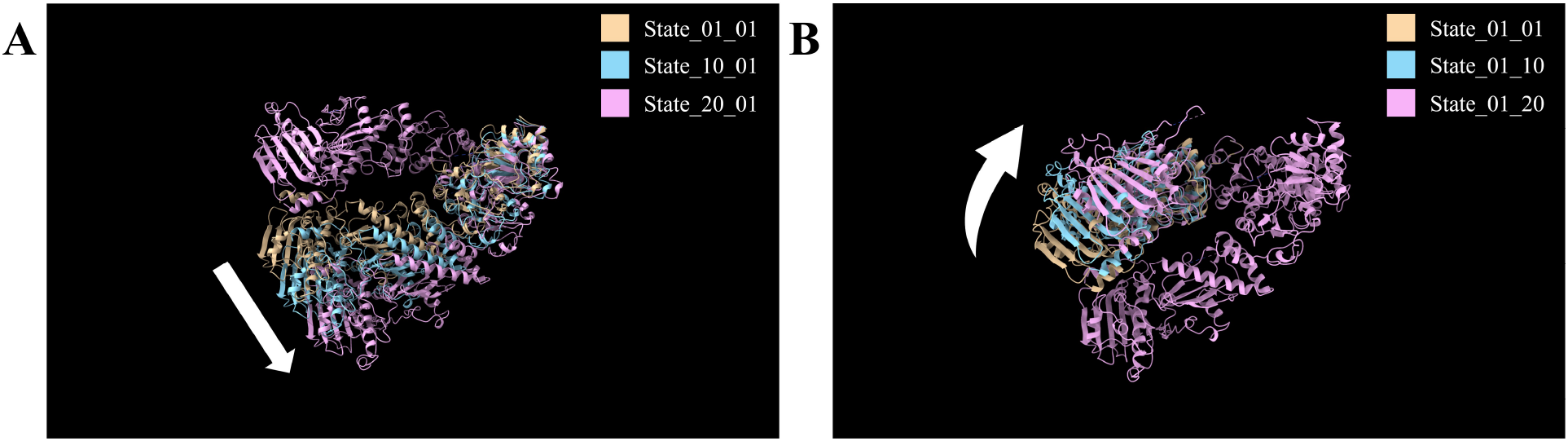
Illustration of conformational changes in the Hsp90 experiment. **A**: Conformational changes in the first direction are modeled by a downward movement of one arm, with a shift of 1 degree for each conformational state. **B**: Conformational changes in the second direction are modeled by an upward rotation of a different arm, with an angular change of 2 degrees for each conformational state.

**Supplementary Figure 5:**
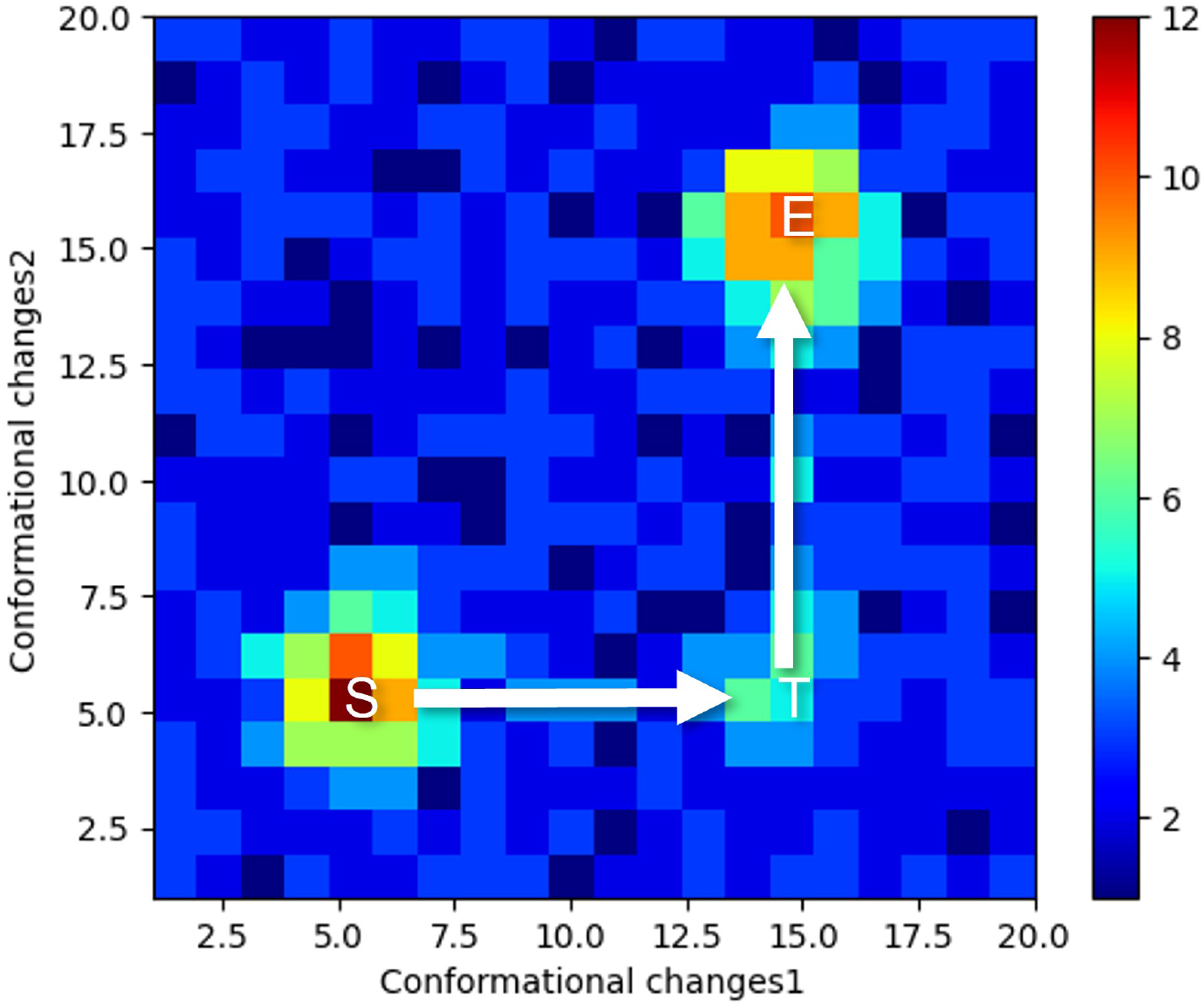
Occupancy map for the Hsp90 experiment. The group at the lower-left corner and the group at the upper-right corner represent the starting (S) and ending (E) conformational states, respectively. The group located at the lower-right corner corresponds to the main transition state (T) we aim to uncover. The occupancy in the background regions is set to 1, and we’ve randomly added 400 points across the entire map. Paths have been included to provide a clear visual guide for the transition processes.

**Supplementary Figure 6:**
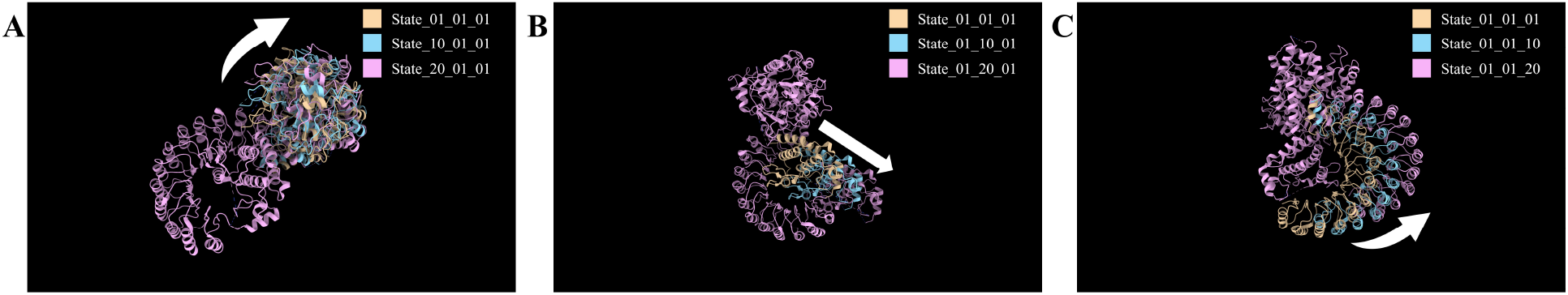
Illustration of conformational changes in the NLRP3 experiment. **A**: The first direction of conformational changes is demonstrated, where the head of the NLRP3 molecule is rotated upward by 2 degrees for each conformational state. **B**: The second direction of conformational changes involves the movement of the binding site by 2 degrees for each conformational state. **C**: The third direction of conformational changes consists of moving the arm upward in the NLRP3 molecule by 2 degrees for each conformational state.

**Supplementary Figure 7:**
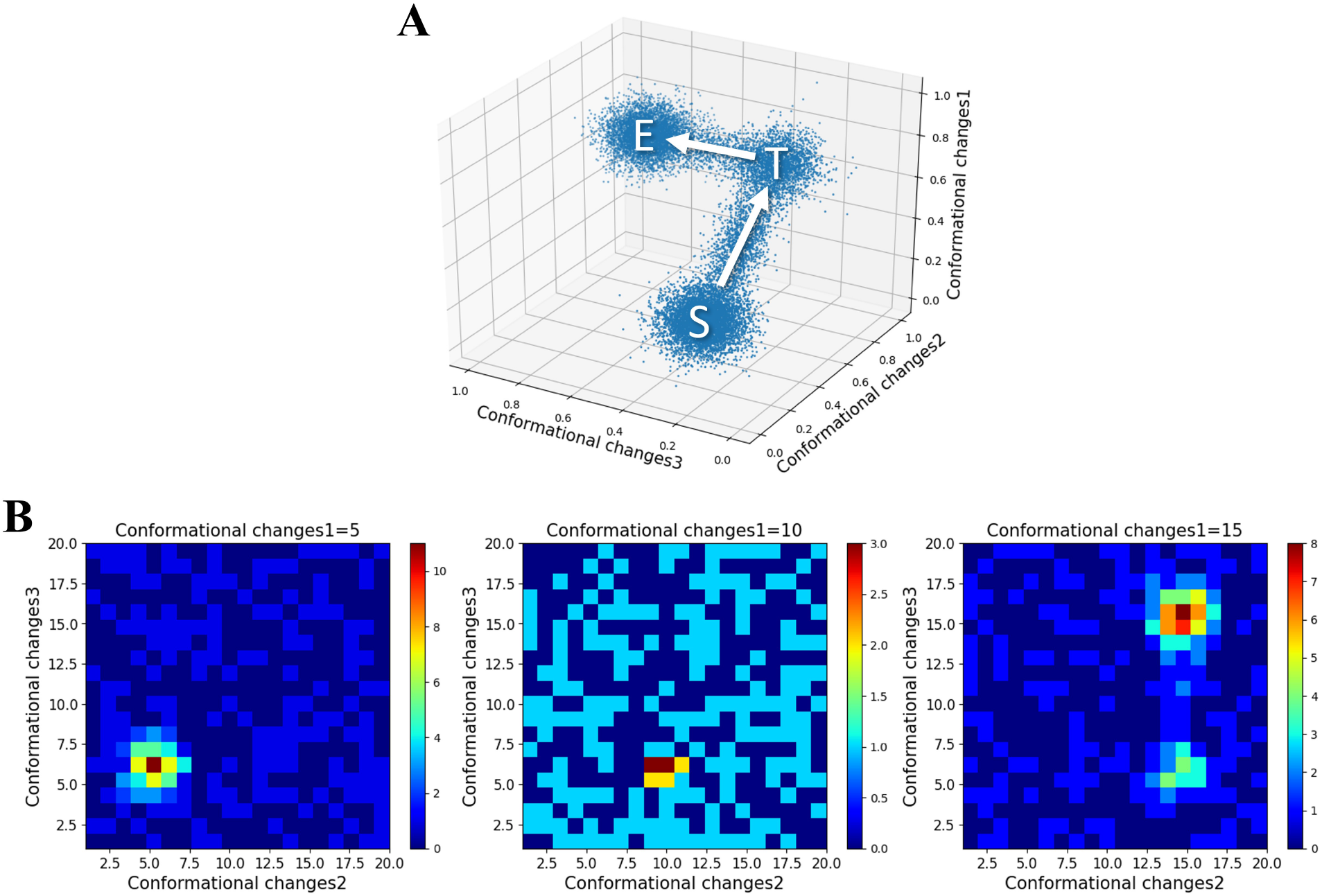
Occupancy map for NLRP3 experiment. **A**: Data points are generated in a 3D space to form the occupancy map, utilizing the same design principle as in the Hsp90 experiment. The two corners denote the starting (S) and ending (E) states, respectively, with the primary transition state (T) situated at another corner. A path links these three groups in the occupancy map. **B**: Slices of the occupancy map along the axis of conformational change 1, with values equal to 5, 10, and 15. Paths have been included to provide a clear visual guide for the transition processes.

**Supplementary Figure 8:**
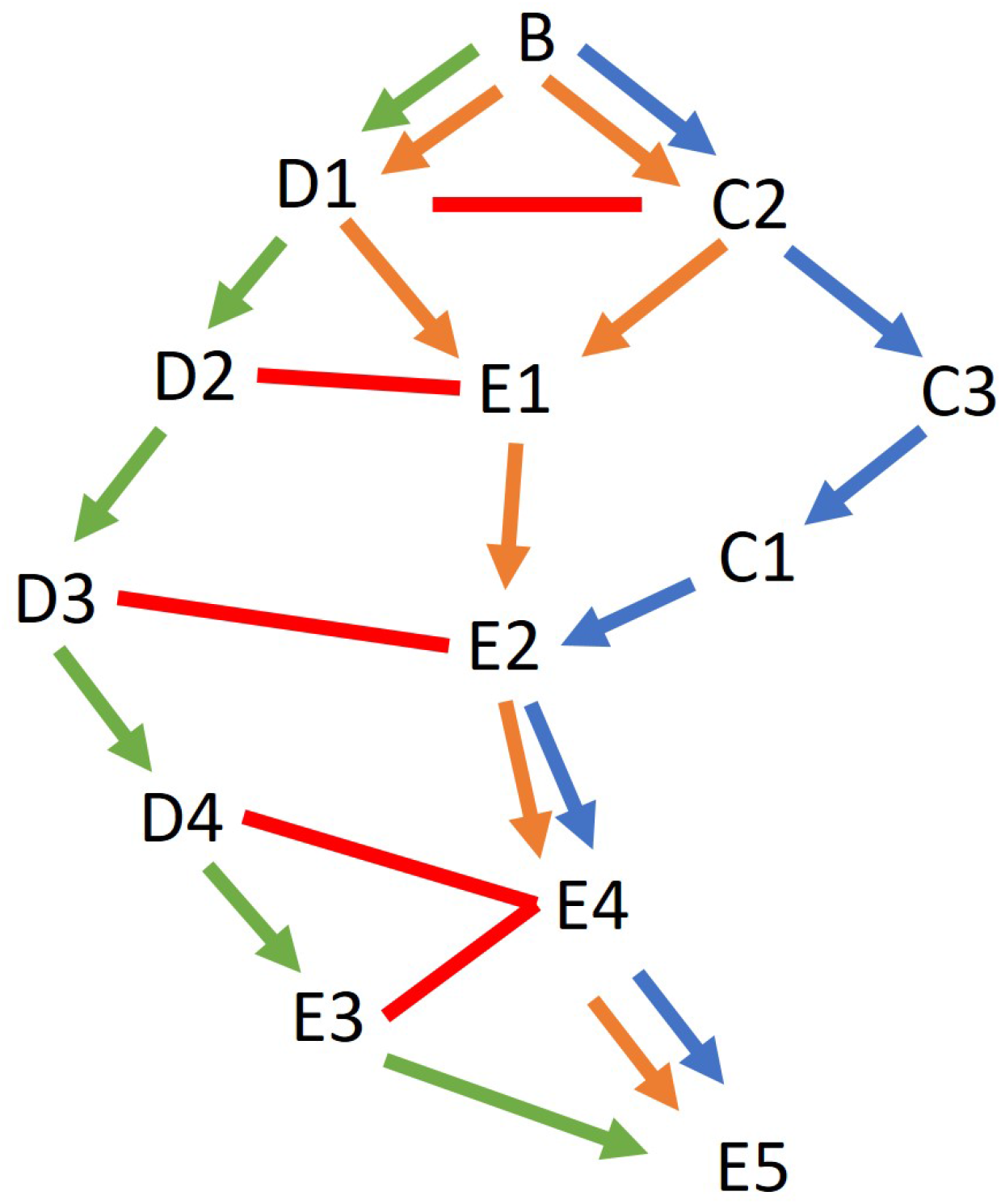
The three conformation change paths suggested in the prior research are visualized in different colors. The first path, consisting of D1, D2, D3, D4, E3, is marked in green. The second path, formed by D1 (or C2), E1, E2, E4, is represented in orange. The third path, incorporating C2, C3, C1, E2, E4, is depicted in blue. Unfavorable transitions are indicated by red lines.

**Supplementary Figure 9:**
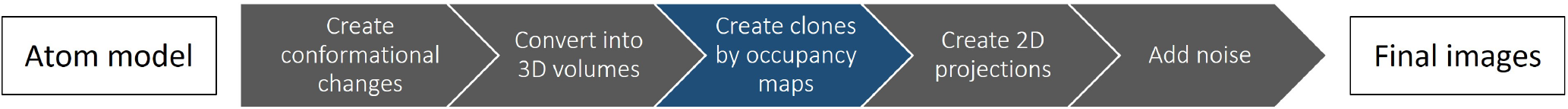
Flowchart of the workflow for the generation of the synthetic dataset with continuous conformational changes

**Supplementary Figure 10:**
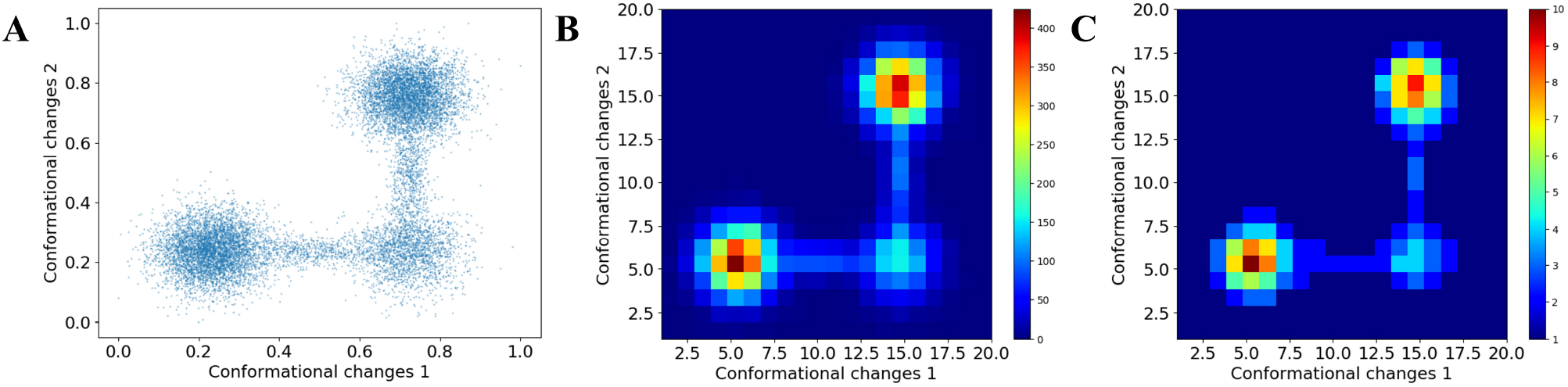
Illustration of the occupancy map generation process with a 2-degree conformational change and 20 conformational states for each coordinate. **A**: Data points are generated using a Gaussian mixture model in a 2D space, embodying the conformational changes with two degrees of freedom. The data point distribution is designed to emulate the presence of different conformational states in the dataset. **B**: Upon segmenting the coordinates into 20 bins, the number of data points within each grid is counted. These counts denote the frequency of each conformational state, with a higher count signifying a more stable or commonly observed state. **C**: To optimize computational efficiency, the counts are rescaled to the desired occupancies, representing the number of clones for each conformational state.

**Supplementary Figure 11:**
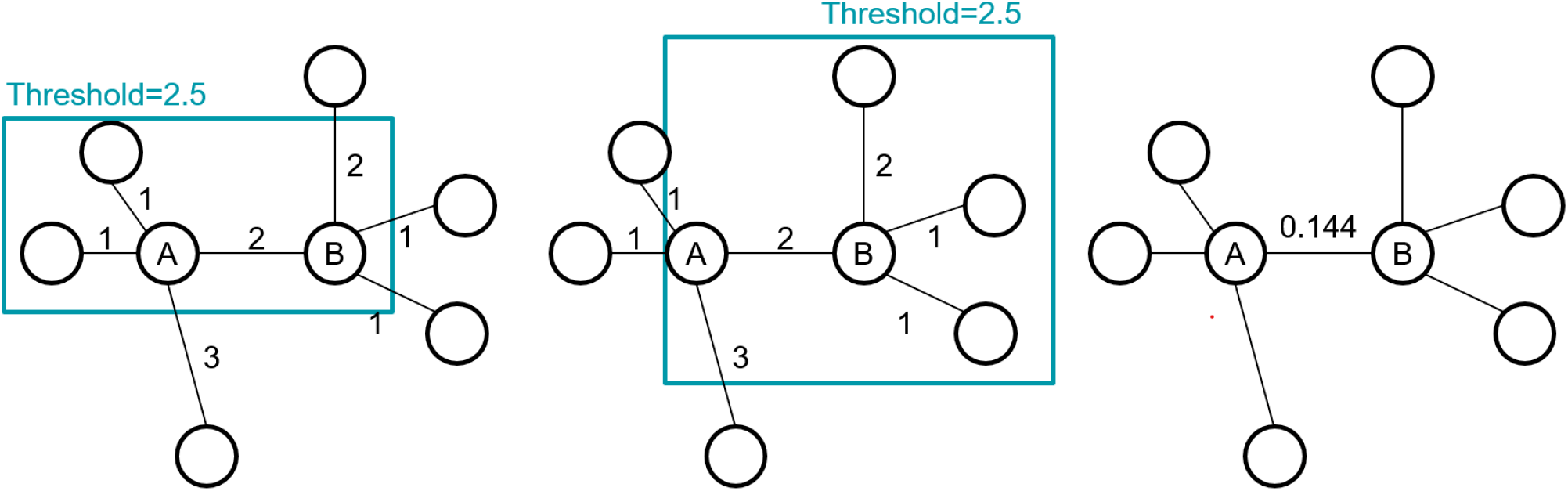
Example of energy edge weighting scheme for a given threshold. The example graph is constructed using 4 nearest neighbors. A predetermined threshold of 2.5 prunes the graph, and node *A* retains 3 neighbors within this threshold. The energy for node *A* is computed as 0.288 using Equation (11). Node *B* maintains 4 neighbors within the threshold, representing the graph’s highest occupancy, thus resulting in an energy of 0. Finally, the edge weights between nodes *A* and *B* are computed as the two nodes’ average energy, which is 0.144.

**Supplementary Figure 12:**
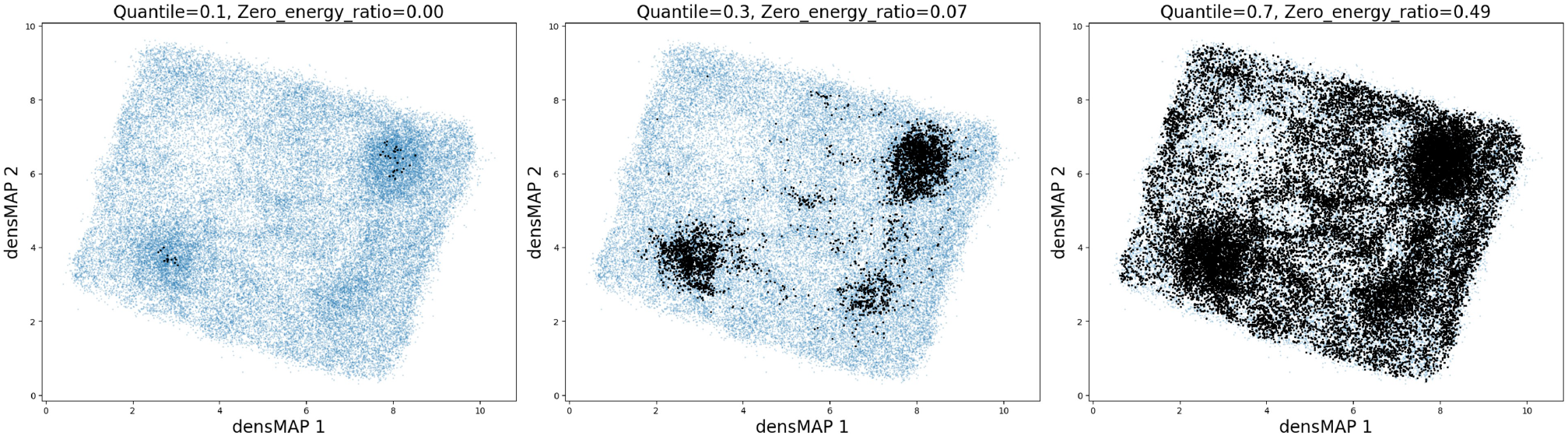
2D visualization of the graph using various quantiles as thresholds in our energy-aware pathfinding algorithm, with threshold values increasing from left to right. The black points denote zero-energy points. As the threshold (expressed in terms of quantiles) increases, the number of zero-energy points within the landscape also increases. Notably, the pattern of these zero-energy points aligns closely with the low-energy region illustrated in Supplementary Figure 1.

**Supplementary Figure 13:**
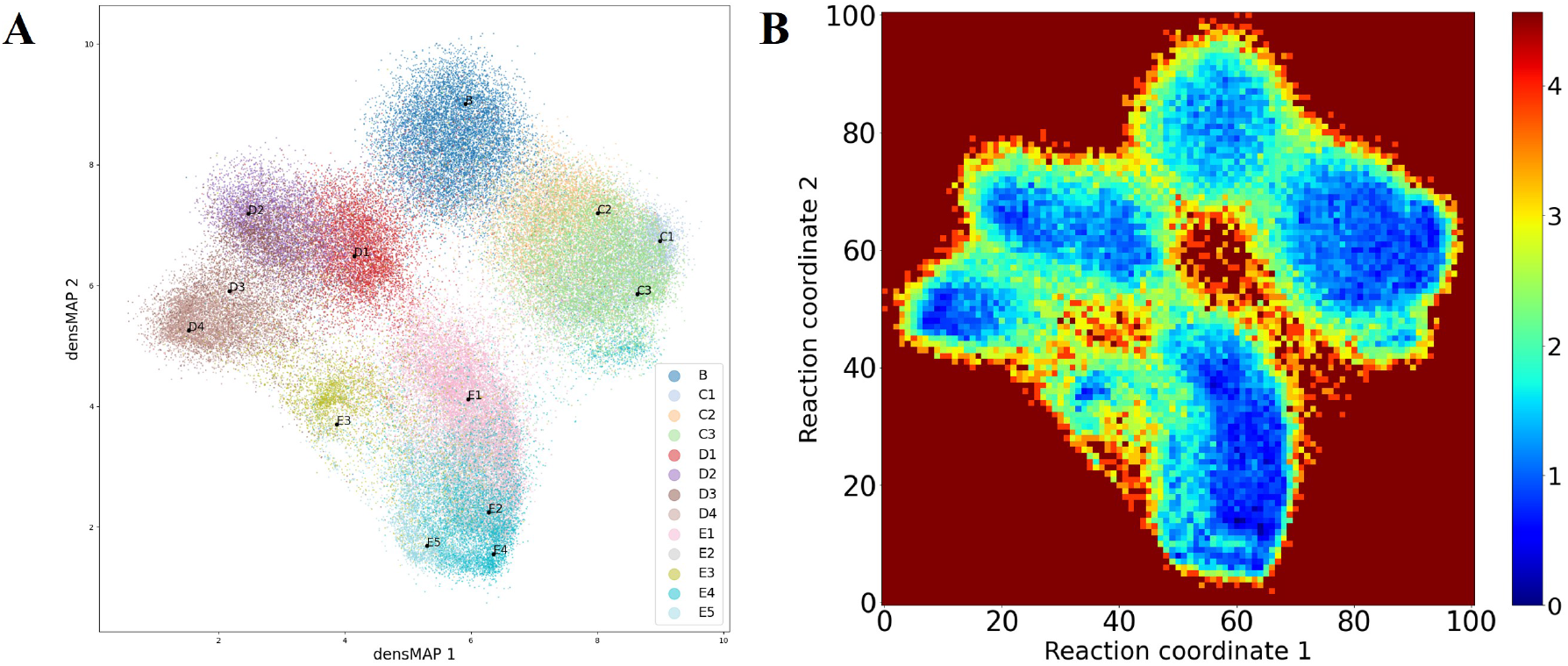
Visualization of the 2D embedding of the latent space and corresponding energy landscape for the EMPIAR-10076 dataset. **A** 2D densMAP visualization of the latent space for the EMPIAR-10076 dataset. Each color represents a distinct conformational state, with the median point in the latent space for each state indicated by black dots. The states labeled A, F, and Discarded are omitted from this visualization. Furthermore, the C4 state, as identified by the cryoDRGN team, is also excluded. **B** The resulting 2D energy landscape derived from applying a 2D histogram to the embedding in **A**. Each coordinate is segmented into 100 bins for occupancy count, with energy estimation calculated using Equation (11).

**Supplementary Figure 14:**
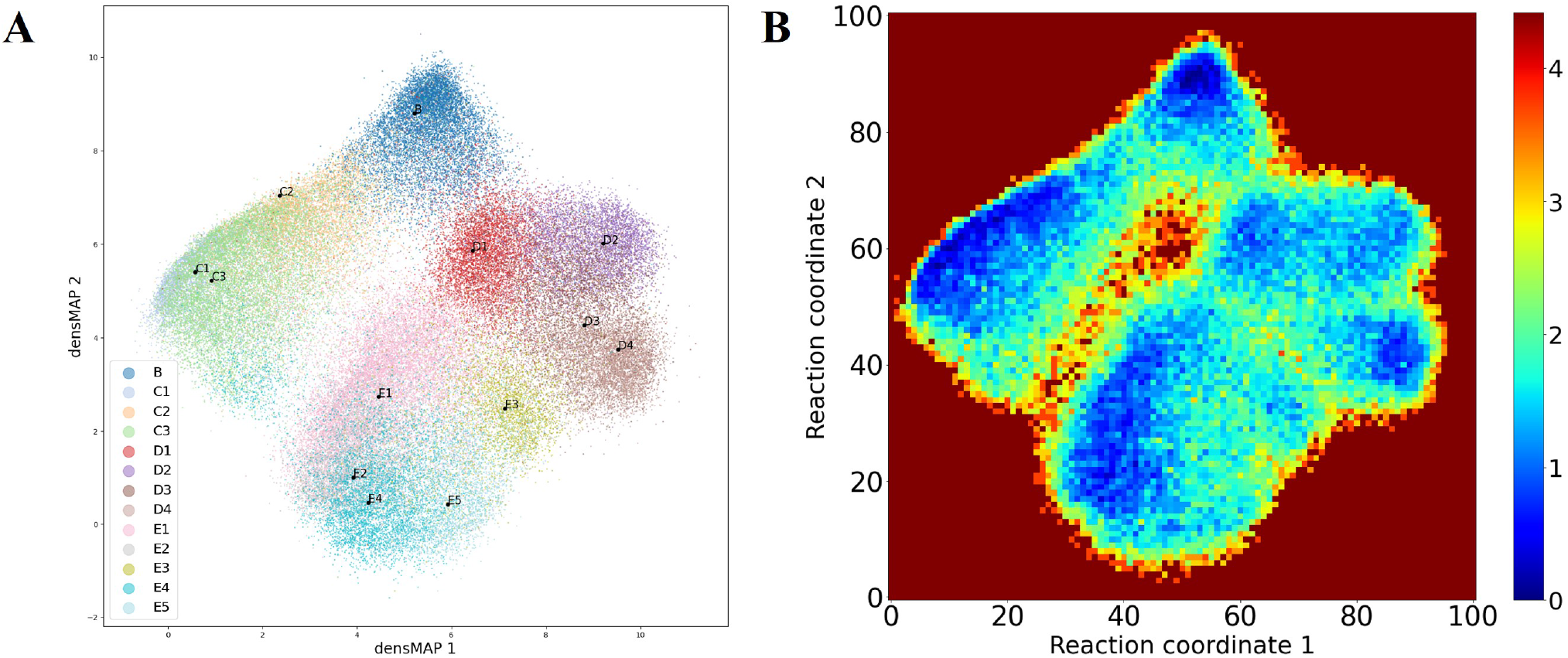
Visualization of the 2D embedding of the latent space for the EMPIAR-10076 Dataset after the analyze-landscape pipeline and corresponding energy landscape. **A**: 2D visualization of the transformed latent space after the analyze-landscape pipeline, using densMAP. Each color represents a unique conformational state, with black dots indicating the median point in this new space for each state. The states labeled A, F, and Discarded are omitted from this visualization. **B**: The derived 2D energy landscape, resulting from the application of a 2D histogram to the embedding in **A**. Each coordinate is divided into 100 bins for occupancy counting, with energy estimation computed using Equation (11).

### Algorithm 1: Proposed energy-aware pathfinding algorithm

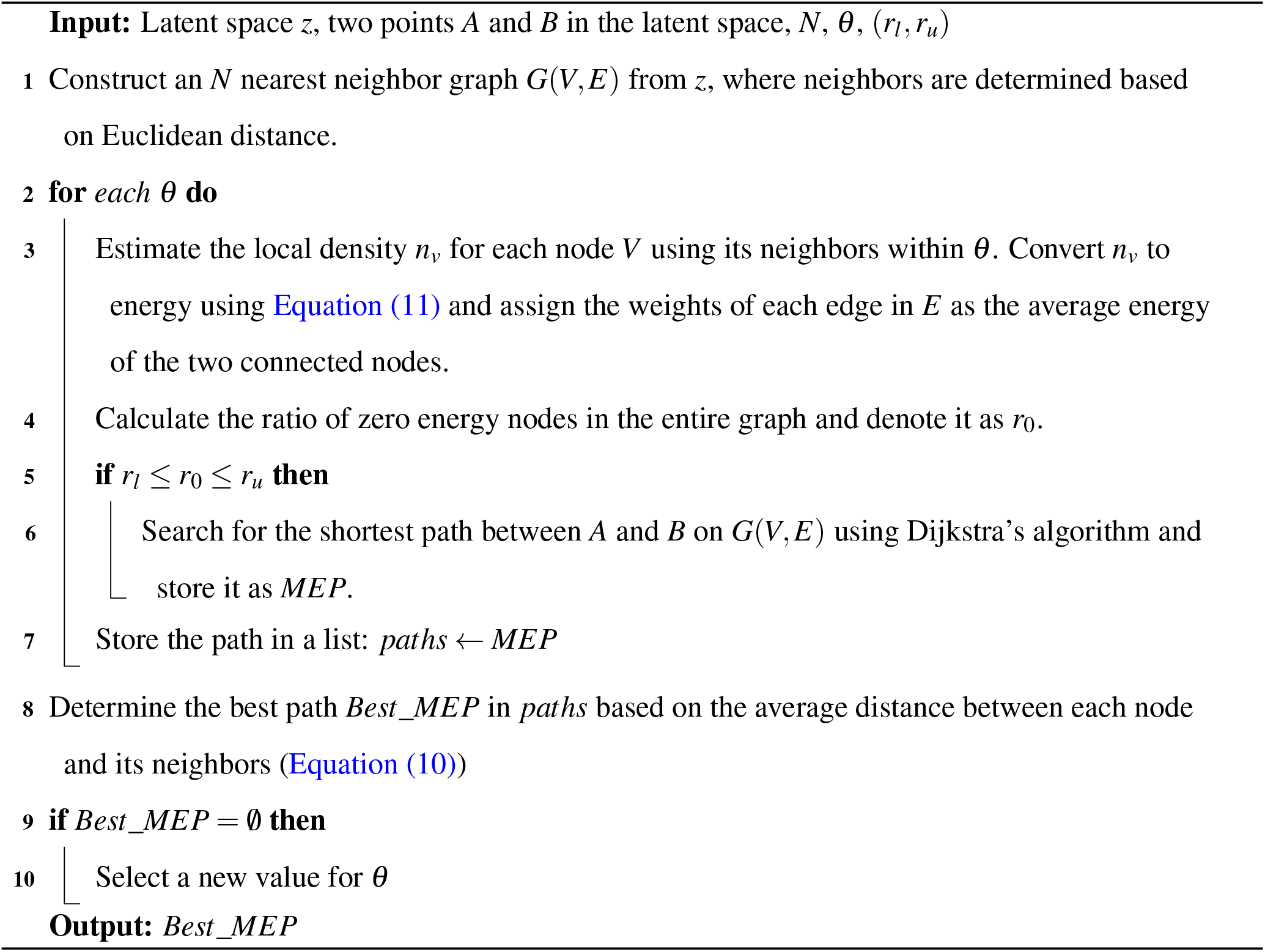

## Footnotes

1 In this study, we consider the conformational landscape as an approximation to the energy landscape.

2 In graph traversal, `max_neighbors` refers to the number of neighbors utilized in constructing a nearest neighborhood graph. Meanwhile, `avg_neighbors` determines the threshold for graph pruning, with a range of (1, `max_neighbors`). The default parameters are set as `max_neighbor = 10` and `avg_neighbors = 5`.

3 At present, the Manifold-EM software requires significant human intervention for hyperparameter tuning and does not support adjusting the dimensionality of the latent space to higher dimensionality.

